# HIV-1 Viremia Not Suppressible By Antiretroviral Therapy Can Originate from Large T-Cell Clones Producing Infectious Virus

**DOI:** 10.1101/2020.01.30.924159

**Authors:** Elias K. Halvas, Kevin W. Joseph, Leah D. Brandt, Shuang Guo, Michele D. Sobolewski, Jana L. Jacobs, Camille Tumiotto, John K. Bui, Joshua C. Cyktor, Brandon F. Keele, Gene D. Morse, Michael J. Bale, Mary F. Kearney, John M. Coffin, Jason W. Rausch, Xiaolin Wu, Stephen H. Hughes, John W. Mellors

## Abstract

**BACKGROUND:** HIV-1 viremia that is not suppressed by combination antiretroviral therapy (ART) is generally attributed to incomplete medication adherence and/or drug resistance. We evaluated individuals referred for non-suppressible viremia (plasma HIV-1 RNA above 40 copies/ml) who reported adherence to ART and did not show drug resistance to their current regimen.

**METHODS:** Samples were collected from at least two time points from eight donors who had non-suppressible viremia for more than six months on ART. Single templates of HIV-1 RNA obtained from plasma and viral outgrowth of cultured cells and from proviral DNA were PCR-amplified and sequenced for evidence of clones of cells that produced infectious viruses. Clones were identified by host-proviral integration site analysis.

**RESULTS:** HIV-1 genomic RNAs with identical sequences were identified in plasma samples from all eight donors. The identical viral RNA sequences did not change over time and lacked resistance to the ART regimen. In four of the donors, viral RNA sequences obtained from plasma matched those sequences from viral outgrowth cultures, indicating that the viruses were replication-competent. Integration sites for infectious proviruses from those four donors were mapped to introns of the *MATR3*, *ZNF268*, *ZNF721/ABCA11P*, and *ABCA11P* genes. The sizes of the clones were from 50 million to 350 million cells.

**CONCLUSION:** Clones of HIV-1-infected cells producing virus can cause failure of ART to suppress viremia despite medication adherence and absence of drug resistance. The mechanisms involved in clonal expansion and persistence need to be defined to eliminate viremia and the HIV-1 reservoir.

## Introduction

In most individuals, combination antiretroviral therapy (ART) suppresses HIV-1 viremia, measured as plasma HIV-1 RNA, to below the limit of detection of commercial assays (20-40 copies/ml), and confers major clinical benefits (1). Despite clinically-effective ART, infected cells persist for the life of the individual and a small subset of the infected cells carry intact proviruses capable of producing infectious virus that can fuel viral rebound when ART is stopped (2–6). Recent work has shown that this reservoir of HIV-1 is sustained by the proliferation of clones of cells with intact proviruses (7–13). Although most of the proviruses in the HIV-1 reservoir remain latent, a small fraction are transcriptionally active (14–17) leading to very low-levels of viremia (usually about 1-3 HIV-1 RNA copies/ml of plasma) that can be detected only with sensitive research assays (18–20). We previously reported that one donor with advanced malignancy had clinically-detected levels of virus in blood (>40 copies/ml) on ART that was comprised of a mixture of drug-resistant virus and wild-type virus, the latter produced by a large clone of cells carrying an intact, infectious provirus (10). Similar cases such as this have not been published.

Current guidelines for treatment of HIV-1 recommend sustained suppression of viremia to below the limit of detection of FDA-approved plasma HIV-1 RNA assays (less than 20-40 copies/ml depending upon the assay). HIV-1 care providers become concerned when viremia is above this level, particularly when it is sustained. Medication review, tests for the presence of HIV-1 drug resistance, and adherence counseling are the standard management approaches used in such cases. Some providers may change or intensify ART regimens by adding additional antiretrovirals in an attempt to achieve greater viremia suppression, although this approach has been ineffective in decreasing low-level viremia in several clinical trials (21–23). We were asked by local care providers to investigate the cause of persistent viremia above the limit of detection of FDA-approved HIV-1 RNA assays in individuals who reported consistent adherence to their medication. We found that the non-suppressible viremia could originate from large clones of infected cells rather than from ongoing viral replication resulting from medication non-adherence or HIV-1 drug resistance.

## Methods

Plasma and peripheral blood mononuclear cells (PBMC) were obtained, following written informed consent, from patients with non-suppressible viremia while on ART under an appropriate human research protocol approved by the University of Pittsburgh’s institutional review board. Non-suppressible viremia was defined as plasma HIV-1 RNA >40 copies/ml for more than six months on ART. Single genome sequencing (SGS) was performed on HIV-1 genomic RNA from plasma and quantitative viral outgrowth assays (qVOA) and on proviral DNA in PBMC (11, 24, 25). Integration sites were obtained using two methods (7, 26). A more detailed description of methods used is provided in the Supplementary Appendix.

## Results

### Study Participants and Characteristics

The characteristics of the eight study participants are summarized in Table 1. All individuals were on long-term suppressive ART (median = 11 years; range 2-18 years) prior to exhibiting non-suppressible viremia for > 6 months (median = 4.1 years; range 1.3-7.4 years). Median pre-ART plasma HIV-1 RNA was 172,413 copies per ml (range 30,375 – 16,700,00) and median nadir CD4^+^ T-cell count was 212 cells/mm^3^ (range 10-314), consistent with substantial immunodeficiency at the time of ART initiation. Other virologic and immunologic characteristics of the donors are provided in Table 2. At the time of referral, the median plasma HIV-1 RNA was 87 copies/mL (range 43-197). Median HIV-1 DNA and cell-associated HIV-1 RNA levels per 1 million PBMC were 1,458 copies (range 373-2,505) and 261 copies (range 29-1,162), respectively (27). The median infectious units per million CD4^+^ T-cells (IUPM) was 0.5 (range 0.1-18.1) (28, 29). The median average pairwise distances (APD) of all proviral sequences was 1.95% (range 0.5%-2.4%), consistent with the diverse HIV-1 populations that arise after long-term infection (Table S1). Coreceptor tropism analyses (Geno2Pheno) of *env* sequences derived from near-full length (NFL) proviral or viral NFL qVOA RNA sequences indicated that viruses were CCR5-tropic (Table S1). Immunophenotyping of PBMC showed no major differences between donors and healthy HIV-1 negative controls in the frequencies of T-cells, B cells, or NK cells or in expression levels of surface activation markers (CD25, CD69, CD38, CD107a), with the exception of higher HLA-DR expression (Table S3), which is consistent with prior observations in HIV-1 positive individuals on effective ART (30).

**Table 1.**
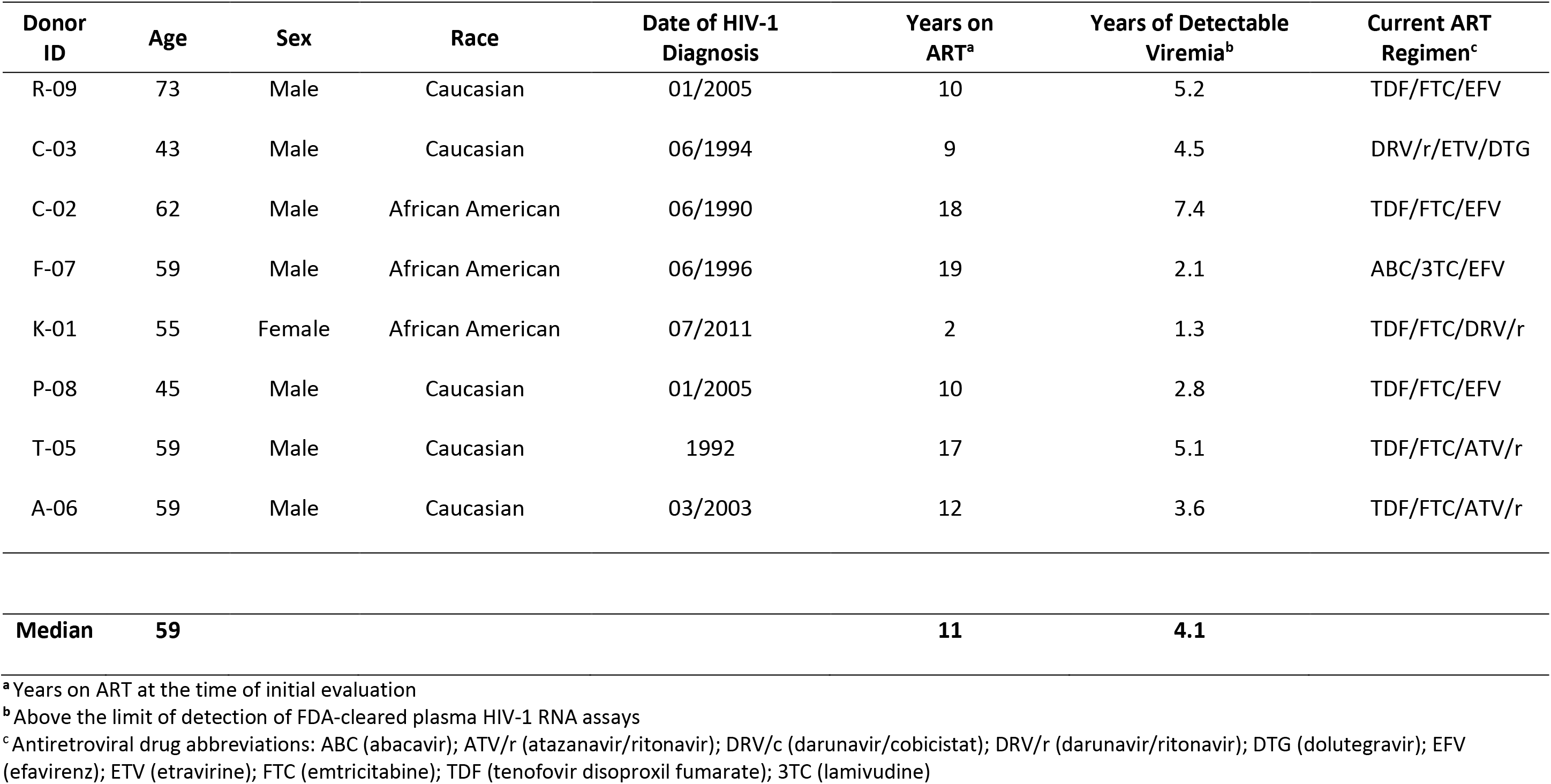
Demographics and Clinical Characteristics of Donors Evaluated for Non-Suppressible Viremia on Antiretroviral Therapy (ART)

**Table 2.** Immunologic and Virologic Characteristics of Donors Referred for Non-Suppressible Viremia

| Donor ID | Pre-ART Plasma HIV-1 RNA (cps/mL) | Nadir CD4<sup>+</sup> T-Cells (cells/mm<sup>3</sup>) | Current CD4<sup>+</sup> T-Cells (cells/mm<sup>3</sup>) | Plasma HIV-1 RNA at Referral (cps/mL)<sup>a</sup> | HIV-1 DNA (cps/10<sup>6</sup> PBMC)<sup>b</sup> | Cell-Associated HIV-1 RNA (cps/10<sup>6</sup> PBMC)<sup>b</sup> | IUPM<sup>c</sup> |
| --- | --- | --- | --- | --- | --- | --- | --- |
| R-09 | 97,000 | 105 | 380 | 197 | 1533 | 139 | 18.1 |
| C-03 | 16,700,000 | 10 | 416 | 62 | 2505 | 1162 | 1.4 |
| C-02 | 30,375 | 286 | 1022 | 184 | 373 | 29 | 0.1 |
| F-07 | 117,068 | 314 | 1023 | 52 | 1603 | 1112 | 3.8 |
| K-01 | 147,189 | 133 | 533 | 68 | 1383 | 74 | 0.6 |
| P-08 | 604,000 | 172 | 444 | 106 | 1056 | 382 | 0.4 |
| T-05 | 197,826 | 299 | 1105 | 113 | 650 | 109 | 0.4 |
| A-06 | 1,877,100 | 251 | 831 | 43 | 1825 | 630 | 0.2 |
| **Median** | **172,413** | **212** | **682** | **87** | **1458** | **261** | **0.5** |

<sup>a</sup> Plasma HIV-1 RNA copies/mL at time of referral determined by FDA-approved Roche CAP/CTM v2.0 or Abbott M2000 platforms
<sup>b</sup> HIV-1 DNA and cell-associated RNA copies/million peripheral blood mononuclear cells (PBMC) measured by quantitative polymerase chain reaction (27)
<sup>c</sup> Infectious units per million (IUPM) total CD4<sup>+</sup> T-cells by quantitative viral outgrowth assay (qVOA) (28, 29)

### Identical Plasma-Derived HIV-1 Sequences Contribute to Detectable Viremia

In all eight donors, groups of identical HIV-1 RNA *gag-propol* sequences derived from single HIV-1 RNA templates (single genome sequencing [SGS]) were found in plasma samples, suggesting but not proving that the identical viruses were derived from clones of infected cells (Figures 1, S1A, S2-S4) (24). The identical RNA sequences comprised a substantial proportion of the total viral sequences in the plasma in all donors (median = 58.6%, range 37.5%-100%) (Table 3). Longitudinal plasma sampling showed no changes in the identical viral sequences over a median sampling duration of 0.8 years (range 0.2-1.2 years). Drug susceptibilities of the plasma viruses, predicted by sequence analysis, showed no evidence of drug resistance to each donor’s current ART regimen, except for a group of identical plasma-derived sequences from donor F-07 that predicted low-level resistance (D67N/K70R/K219Q) to only abacavir comprising 53.8% of the total plasma sequences (Table 3). Drug concentrations for all donors were within the expected therapeutic range based upon target trough concentrations (Table S2).

**Table 3.** Single Genome Sequence Matches Between Plasma HIV-1 RNA, Proviruses, and Viral Outgrowth Cultures

| Donor ID | % Identical Plasma HIV-1 RNA Sequences<sup>a</sup> | % Proviral Sequences Matching Dominant Plasma HIV-1 RNA Sequences (% matching)<sup>b</sup> | % of qVOA p24<sup>+</sup> Wells with Matching Sequences in Plasma HIV-1 RNA<sup>c, d</sup> | % Proviral Sequences Matching Dominant qVOA HIV-1 RNA Sequences (% matching)<sup>e</sup> |
| --- | --- | --- | --- | --- |
| R-09 | 50.0% | 10.7% | 100% (10/10) | 10.7% |
| C-03 | 63.3% | 2.9% | 66.7% (2/3) | 2.9% |
| C-02 | 90.7% | 7.5% | 100% (3/3) | 7.5% |
| F-07 | 53.8% | 0.3% | 0% (0/4) | 3.0% |
| K-01 | 37.5% | 3.9% | 0% (0/4) | <3.9% |
| P-08 | 100% | 8.0% | 0% (0/3) | <8.0% |
| T-05 | 76.7% | <0.7% | 0% (0/3) | <0.7% |
| A-06 | 48.4% | <0.7% | 0% (0/2) | <0.7% |

<sup>a</sup> Percent (%) of all HIV-1 RNA single genome sequences from plasma (1.3 kb of *gag-pro-pol*) that are identical or differ by 1-2 bases from the group of identical sequences
<sup>b</sup> Identical proviruses (single genome sequences) in PBMC that match HIV-1 RNA sequences from plasma (1.3 kb of *gag-pro-pol*)
<sup>c</sup> Quantitative Viral Outgrowth (qVOA) HIV-1 RNA single genome sequences (1.3 kb of *gag-pro-pol*)
<sup>d</sup> Percent (%) and number of *gag* p24 antigen positive (p24<sup>+</sup>) qVOA wells with identical sequences that match plasma HIV-1 RNA sequences (1.3 kb of *gag-pro-pol*)
<sup>e</sup> Percent (%) of total proviral HIV-1 DNA single genome sequences that match identical qVOA HIV-1 RNA single genome sequences (1.3 kb of *gag-pro-pol*)
**Bold** indicates that the proviruses in clonally-expanded cells are intact (replication-competent) and either contribute to persistent infectious viremia (CF-02, CN-03, RF-09) or can be induced ex vivo (FN-07) to produce infectious virus (qVOA)

**Figure 1.**
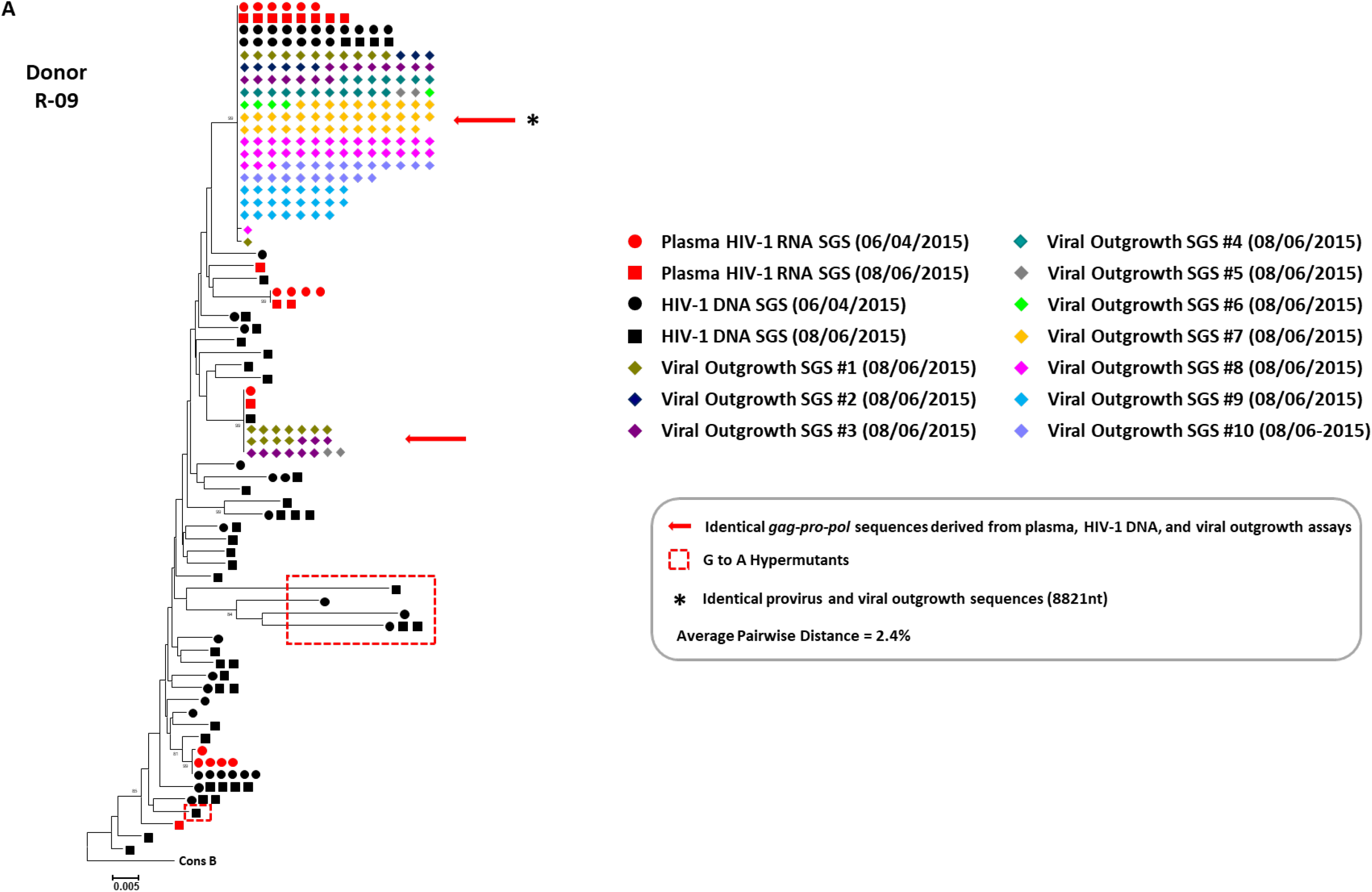
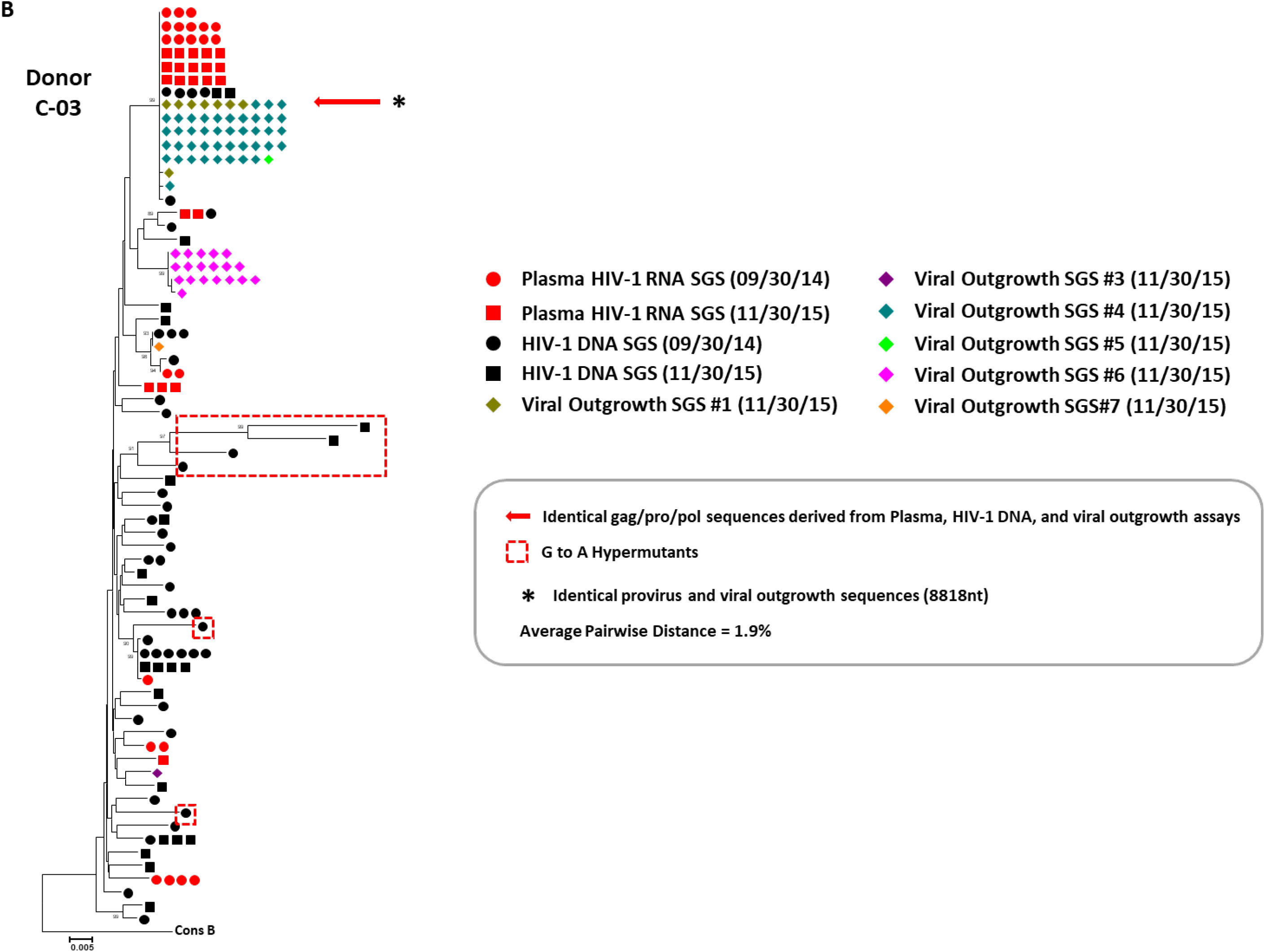
Representative Neighbor-Joining *p*-Distance Phylogenetic Trees of Plasma HIV-1 RNA-, HIV-1 DNA-, and Quantitative Viral Outgrowth Assay (qVOA)-Derived Sequences from Donors R-09 and C-03 with Intact Replication-Competent Proviruses Producing Non-Suppressible Viremia. The trees were rooted to a subtype B consensus sequence. Single genome sequences (SGS) of a portion of *gag (p6)*, all of *pro*, and the portion of *pol* encoding the 1^st^ 300 amino acids of reverse transcriptase (*gag-pro-pol*) were obtained from plasma HIV-1 RNA, HIV-1 DNA in peripheral blood mononuclear cells, and culture supernatants from p24^+^ qVOA wells for donor R-09 (Panel A) and donor C-03 (Panel B) (24). Red circles and squares represent plasma-derived sequences from two time points. Black circles and squares represent HIV-1 DNA-derived sequences from two time points. Different colored diamonds represent viral outgrowth assay-derived sequences from independent p24^+^ quantitative viral outgrowth assay (qVOA) wells. A red arrow shows identical *gag-pro-pol* sequences from plasma HIV-1 RNA-, HIV-1 DNA-, and p24^+^ qVOA HIV-1 RNA-derived sequences. The asterisk shows matching sequences of provirus (near-full length HIV-1 DNA and host to full length provirus to host amplicons) and near-full length viral RNA sequences from p24^+^ wells. HIV-1 DNA sequences with G to A hypermutations are enclosed in red hashed boxes. The viral outgrowth sequence variants that differ by 1-2 nucleotides can be attributed to either ex vivo replication or errors introduced during cDNA synthesis. Average pairwise distances (APD) calculated by MEGA v6.0 using HIV-1 DNA sequences excluding hyper-mutated sequences.

### Identical Plasma-, Proviral-, and Viral Outgrowth-Derived HIV-1 Sequences

Genomic DNA (gDNA) from PBMC or purified CD4^+^ T-cells and HIV-1 RNA from p24^+^ qVOA wells were analyzed by *gag-pro-pol* single genome sequencing (SGS) (Figure S1B-S1D). Matching Identical *gag-pro-pol* sequences were found in proviral DNA and plasma HIV-1 RNA in six of the eight donors (K-01, C-02, C-03, F-07, P-08, and R-09) (Figures 1, S2-S4) (24). In three of these donors (C-02, C-03, and R-09), the identical viral sequences in plasma also matched multiple qVOA-derived sequences from p24^+^ wells (Table 3, Figures 1, S2). In one donor (F-07), there were identical proviral and qVOA-derived sequences, but no sequences were found in plasma that matched (0 of 39). A different provirus from this donor matched plasma viral sequences, but these sequences were not found in p24^+^ qVOA wells (Table 3, Figure S3).

### Persistent Viremia from Intact, Replication-Competent HIV-1 Proviruses

Near-full length viral sequence analyses were performed on samples from the four donors for which there were identical *gag-pro-pol* sequence matches between plasma and p24^+^ qVOA cultures. Phylogenetic analyses of sequences from donor R-09 suggested strongly that the persistent viremia was being produced by a clone of cells that carried an intact, replication competent provirus: six NFL HIV-1 DNA and four NFL HIV-1 RNA sequences from p24^+^ qVOA wells were identical over 8821nt and matched the *gag-pro-pol* sequences found in plasma in longitudinal samples obtained 2 months apart (Figure 1A and Table 3). Similarly, phylogenetic evidence for a clonal origin of persistent viremia was also obtained for donors C-03 and C-02 from whom identical NFL proviral and qVOA-derived sequences (8818 and 8906nt, respectively) were obtained that matched plasma-derived *gag-pro-pol* sequences (Figures 1B and S2, Table 3). Identical viral RNA sequences were obtained from the plasma of these two donors over periods of 14 and 10 months, respectively. In donor F-07, one NFL proviral sequence (8819nt) matched identical *gag-pro-pol* plasma-derived sequences from two longitudinal samples taken 13 months apart, but matching viral sequences were not found in p24^+^ qVOA wells (Figure S3, Table 3). Identical NFL sequences (8804nt) of a different provirus were found in blood that matched the qVOA-derived *gag-pro-pol* sequences, but as noted above, no matches were found in the viral RNA in plasma. Collectively, these analyses suggest that there are clones of cells in the blood of these four donors that can carry replication-competent proviruses, which we have termed repliclones.

### Proof That Non-Suppressible Viremia can Originate from Clones of Infected Cells

We performed integration site analysis (ISA) (Figure S1E) on genomic DNA extracted from purified CD4^+^ T-cells from the donors (R-09, C-03, C-02, and F-07) with plasma and/or proviral sequences that matched viruses obtained from qVOAs. In samples from each of these four donors, we found multiple identical integration sites for the proviruses whose sequences matched the identical sequences in plasma and qVOA. Table 4 and Figure S5 show the locations and orientations of proviruses in the host genome (31, 32). All integration sites identified were within introns of known host genes including ABCA11P, MATR3, ZNF268, and ZNF721. The exact location and full sequence of each of the intact proviruses was confirmed by amplifying and sequencing the entire provirus and flanking host sequences directly from genomic DNA extracted from PBMC (Figure S1F). We determined the frequency of these integrated intact proviruses compared to all other proviruses (Table 4) and found them to be a minor fraction of all infected cells (0.03%-1.1%) (7). We also estimated the total size of each of the repliclones within each of the four individuals based upon the approximate sizes of the CD4^+^ T-cell populations in each of the donors and the frequency of the specific, intact provirus per million CD4^+^ T-cells (Table 2) (27, 33). The total size of the repliclones ranged from approximately 50 million to 350 million CD4^+^ T-cells (Table 4). These findings indicate that individual repliclones are large enough to make an important contribution to the HIV-1 reservoir even though they comprise a very small fraction of all infected cells.

**Table 4.** Integration Site Analyses of Intact Replication Competent Proviruses in Expanded Clones

| Donor | Intact Provirus Integration Site<sup>a</sup> | Gene<sup>b</sup> | Location<sup>c</sup> | Proviral Orientation Relative to Gene<sup>d</sup> | Frequency of Clone by Integration Site Assay<sup>e</sup> | Estimated Size of Clone (# of Cells)<sup>f</sup> |
| --- | --- | --- | --- | --- | --- | --- |
| R-09 | Chr4: 425,766 | ABCA11P | Intron | - | 0.03% (3/9939) | $3.5 \times 10^8$ |
| C-03 | Chr12: 133,777,660 | ZNF268 | Intron | - | 1.10% (44/4164) | $1.3 \times 10^8$ |
| C-02 | Chr5: 138,660,320 | MATR3 | Intron | - | 0.65% (3/459) | $5.2 \times 10^7$ |
| F-07 | Chr4: 443,311 | ZNF721/ABCA11P | Intron | - | 0.05% (4/8407) | $8.0 \times 10^7$ |

<sup>a</sup> Chromosomal location of the proviral 3' LTR in the human genome hg19 reference using UCSC Genome Browser (31, 32)
<sup>b</sup> Gene containing the provirus
<sup>c</sup> Intragenic location of provirus
<sup>d</sup> Proviral orientation relative to gene: positive (+) in same orientation as gene; negative (-) in opposite orientation to gene
<sup>e</sup> Frequency of integrant relative to total integrants determined by integration site analyses (7)
<sup>f</sup> Estimated size of each clone derived from the frequency of matching proviral sequences in CD4<sup>+</sup> T-cells frequency and the total number of CD4<sup>+</sup> T-cells in each donor. Total CD4<sup>+</sup> cells calculated from % CD4<sup>+</sup> cells × total body lymphocytes ( $2 \times 10^{12}$ ) (33)

## Discussion

Here we describe a new cause of HIV-1 plasma viremia observed in standard clinical practice that is not suppressible by ART and show that the viremia can arise from large clones of HIV-1 infected cells. Extensive virologic analyses of the eight individuals referred for non-suppressible viremia on ART revealed important insights for patient management and for efforts to cure HIV-1 infection. Although each of the eight individuals have unique features, the consistent finding, in all donors, was that non-suppressible plasma viremia consisted of one or more large groups of identical viral sequences. The largest group of identical viral sequences in plasma comprised 37.5%-100% (median 53%) of all HIV-1 RNA in the plasma (Table 3). Similar findings of identical viral sequences in plasma were previously reported among individuals on effective ART, although the origin of the identical viruses were not identified (34). In the current study, drug resistance to the donors’ ART regimen was not evident by sequence analysis nor was medication non-adherence since antiretroviral drug concentrations measured in random plasma samples were in the therapeutic range. Longitudinal analyses of the viral sequences in all donors showed no evidence of virus evolution over time, indicating that these viruses originated from a stable, non-evolving reservoir of infected cells. The important implications of these findings for clinical management are that changes in the antiretroviral drug regimen or efforts to enhance drug adherence are unlikely to change the viremia that is being produced by infected cell clones, which are not affected by current antiretroviral drugs. Drugs that block virion production from cells that are already infected would be expected to lower viremia but such agents are not currently available. HIV-1 protease inhibitors that are available block virion maturation into infectious particles but do not reduce virion production from cells that are already infected (35). Although the frequency of non-suppressible viremia of clonal origin is unknown, the cases referred to us for evaluation were from a group of approximately 2000 patients cared for at the referring centers.

In three of the eight donors, identical sequences were found in plasma, proviral DNA, and qVOA, indicating that the viruses were replication-competent and potentially transmissible by exposure to blood. In a fourth donor, identical sequences were obtained from proviral DNA and qVOA that did not match any viral sequences in plasma, indicating for that donor, the viruses in the plasma had a different origin. In the remaining four donors, there were viruses with identical sequences found in the plasma, but these sequences did not match any of the sequences obtained by qVOA (Figure S4). In two of these four donors, proviruses were not found with sequences that matched HIV-1 RNA in plasma (T-05, 0 of 114 sequences; A-06, 0 of 77 sequences), suggesting that the virus-producing cells were in lymphoid tissues but not in blood (Figure S4C-D). The lack of matches between the virus in the plasma and the viruses recovered from qVOA is not surprising given that plasma virus could either be non-infectious or not induced in a single qVOA (12). It is also possible that the clones of cells producing the plasma viremia are relatively small and might not be present in blood samples used for qVOA (36).

In the four donors with clones containing intact proviruses that produced infectious virus, we identified their corresponding integration sites (Tables 3-4 and Figure S5). In all cases the integration sites were within introns of known genes and the proviruses were orientated opposite to the host gene, a pattern that is consistent with what is known about HIV-1 integration sites both in cultured cells and in donors (7, 13, 37). Two of the four proviruses were integrated in ABCA11P and ZNF721 within about 17,500 nucleotides of each other, near the end of chromosome 4. Using a larger collection of integration sites obtained from multiple donors on ART, we found no evidence of positive in vivo selection for proviruses in either of the two overlapping genes (ABCA11P/ZNF721) in which the two intact proviruses are integrated (data not shown), thus their integration in the same overlapping gene is likely to be a coincidence.

Integration site analyses showed that the clones carrying intact proviruses were a small fraction (0.03%-1.1%) of all of the infected cells. However, given the large number of CD4^+^ T-cells in the human immune system, the clones were estimated to comprise large numbers of infected cells (50 million-350 million cells) based upon the percentage of matching proviruses and estimated total body CD4^+^ T-cells (Table 4) (33). The very low frequency of cells infected with the specific intact proviruses makes analyses of the CD4^+^ T-cell subsets involved very challenging given the limitations of cell sampling and current technologies to detect specific proviruses, although work is in progress.

Our findings have important implications for efforts to develop a cure for HIV-1 infection. It is widely believed that the latent HIV-1 reservoir is the main obstacle to a cure. Some have questioned the relevance of persistent viremia on ART, but our findings show that some of the viruses that persist in plasma can be infectious and could quickly initiate viral rebound if ART is discontinued (34, 38, 39). It is possible, in cases in which there are much lower levels of viremia, that there are smaller clones of infected cells releasing infectious virus at levels that are hard to detect but are still capable of rekindling viral replication if ART is interrupted. Importantly, the mechanisms that allow large clones to expand and continually produce virus need to be defined. At present, we cannot determine whether there is a small fraction of each repliclone that produces virus continuously, somehow evading killing by the virus or the immune response, or whether the virus producing cells die after their proviruses are expressed, to be replaced by equal numbers of cells that are able to produce virus. The large size of the clones we found that carry replication-competent proviruses suggest that developing therapies to eliminate them or control their ability to produce virus represents a formidable challenge.

This project has been funded in part by the Intramural Research Programs of the National Cancer Institute, National Institutes of Health (NIH), under Contract No. HHSN261200800001E (EKH, KWJ, LDB, SG, MDS, JLJ, CT, JCC, BFK, MJB, MFK, JWR, XW, SHH, JWM) and 75N91019D00024 (SHH); the Howard Hughes Medical Research Fellows Program, Howard Hughes Medical Institute, Bethesda, MD (JKB); the Bill & Melinda Gates Foundation award OPP1115715 (EKH, KWJ, LDB, MDS, JWM); the Office of AIDS Research (MJB, MFK, JWR); The American Cancer Society (JMC) and the National Cancer Institute through a Leidos subcontract, l3XS110 (JMC); and the National Institute for Allergy and Infectious Diseases, NIH, to the I4C Martin Delaney Collaboratory (UM1AI126603; JLJ, JWM), the University of Rochester Center for AIDS Research (P30AI078498; GDM) and the University of Rochester HIV/AIDS Clinical Trials Unit (UM1 AI069511-08; GDM).

The content of this publication does not necessarily reflect the views or policies of the Department of Health and Human Services, nor does mention of trade names, commercial products, or organizations imply endorsement by the U.S. Government.

J.W.M has served as a consultant to Gilead Sciences, Merck Co. Inc., Xi’an Yufan Biotechnologies, and Accelevir Diagnostics, and owns share options in Co-Crystal Pharmaceuticals, Inc. None of the other authors report conflicts of interest.

We thank the individuals who volunteered for this study. We also thank the referring physicians Bernard Macatangay, MD, and Sharon Riddler, MD, from the University of Pittsburgh Medical Center (UPMC), and Chiu-Bin Hsiao, MD, from Allegheny Health Network. We are grateful for the efforts of the clinical staff at the UPMC HIV-AIDS Program, especially Renee Weinman and Jamie Ideluca. We are also thankful for the efforts of Lorraine Pollini for proofreading, formatting, and submission of this manuscript.

## APPENDIX

Authors’ full names and academic degrees are as follows: Elias K. Halvas, Ph.D., Kevin Joseph, B.S., Leah D. Brandt, Ph.D., Shuang Guo, Ph.D., Michele D. Sobolewski, M.S., Jana L. Jacobs, Ph.D., Camille Tumiotto, M.D., Ph.D., John K. Bui, M.D., Joshua C. Cyktor, Ph.D., Brandon F. Keele, Ph.D., Gene D. Morse, Pharm.D., Michael J. Bale, B.S., Mary F. Kearney, Ph.D., John M. Coffin, Ph.D., Jason W. Rausch, Ph.D., Xiaolin Wu, Ph.D., Stephen H. Hughes, Ph.D., and John W. Mellors, M.D.^*^

Authors’ affiliations are as follows: From the Department of Medicine, University of Pittsburgh, Pittsburgh, PA 15213, USA (E.K.H., K.W.J., L.D.B., M.D.S., J.L.J., C.T., J.C.C, J.W.M); New York-Presbyterian Weill Cornell Medical Center, Department of Medicine, New York, USA (J.K.B). AIDS and Cancer Virus Program, Frederick National Laboratory for Cancer Research, Frederick, MD, 21702, USA (B.F.K.); HIV Dynamics and Replication Program, National Cancer Institute, Frederick, MD 21702, USA (M.J.B, M.F.K., S.H.H.); Department of Molecular Biology and Microbiology, Tufts University, Boston, MA 02111, USA (J.M.C.); Basic Research Laboratory, Center for Cancer Research, National Cancer Institute, Frederick, MD 21702, USA (J.W.R.), Leidos Biomedical Research, Inc., Frederick, MD 21702, USA (X. W., S. G.); Translational Pharmacology Research Core, Center for Integrated Global Biomedical Sciences, School of Pharmacy and Pharmaceutical Sciences, University at Buffalo, Buffalo, NY 14221 (G.D.M).

Address reprint requests or correspondence to Dr. Mellors at Division of Infectious Diseases, Department of Medicine, University of Pittsburgh, 3550 Terrace St., Pittsburgh, PA 15261 or at

