## Supplemental Figures and Tables for "HIV-1 Viremia Not Suppressible By Antiretroviral Therapy Can Originate from Large T-Cell Clones Producing Infectious Virus"

This appendix has been provided by the authors to give readers additional information about their work.

**SUPPLEMENTARY APPENDIX**

**For Halvas EK, et al.,**

**HIV-1 Viremia Not Suppressible By Antiretroviral Therapy Can Originate**

**from Large T-Cell Clones Producing Infectious Virus**

**TABLE OF CONTENTS**

| | |
|---|---|
| SUPPLEMENTARY METHODS | 3 |
| SUPPLEMENTARY FIGURES S1-S6 | 9 |
| SUPPLEMENTARY TABLES S1-S5 | 22 |
| REFERENCES | 27 |

### SUPPLEMENTARY METHODS

#### PARTICIPANTS AND SAMPLE COLLECTION

The University of Pittsburgh Institutional Review Board approved the study, and all participants provided written informed consent. Study participants were referred from the University of Pittsburgh Medical Center HIV-AIDS Program or from the Allegheny Health Network Positive Health Clinic and enrolled into the study at the University of Pittsburgh Clinical Trials Unit. Inclusion criteria for the study were: 1) initial suppression of plasma HIV-1 RNA to below the limit of detection of commercial assays (<20 or <40 copies/mL) on a DHHS-recommended ART regimen; 2) followed by clinically-detectable plasma viremia (HIV-1 RNA >40 copies/mL) for > 6 months, as measured by the COBAS Ampliprep/COBAS Taqman, v2.0 assay (TMv2.0) (Roche) or the m2000sp/RealTime HIV-1 Assay (Abbott); and 3) assessment by the referring physician that the patient was fully compliant with their ART regimen. Switches in ART regimen or intensification with another antiretroviral did not exclude participants from the study.

Longitudinal samples were collected at two or more time points as large-volume phlebotomy (100-180mL) or plasmapheresis and leukapheresis. Plasma, peripheral blood mononuclear cells (PBMC), and total $CD4^+$ T-cells were isolated and stored as previously reported (1).

#### OVERVIEW OF APPROACH TO EVALUATING NON-SUPPRESSIBLE VIREMIA

The schematic in Figure S1 provides an overview of the multistep methodological approach to evaluating non-suppressible viremia and confirming its clonal cellular origin. The individual methods used are described below.

**QUANTIFICATION OF PLASMA HIV-1 RNA AND CELLULAR HIV-1 DNA AND RNA**

HIV-1 DNA and RNA in PBMC or total $CD4^+$ T-cells was quantified by qPCR targeting the 3' end of integrase as previously reported (1).

**QUANTITATIVE VIRAL OUTGROWTH ASSAY**

Quantitative viral outgrowth assays (qVOA) were performed using total $CD4^+$ T-cells. The frequency of HIV-1 infected cells carrying an inducible infectious provirus (IUPM) was determined by a maximum likelihood method, as reported (2-4).

**SINGLE GENOME AMPLIFICATION AND SEQUENCING (SGS)**

SGS of *gag* (*p6*), *pro*, and the first 300 amino acids of *pol* (*gag-pro-pol*) was performed using HIV-1 RNA from plasma or from $p24^+$ qVOA wells or from cellular HIV-1 DNA as follows: i) endpoint dilution of extracted nucleic acid to single HIV-1 template per PCR reaction as determined by Poisson distribution statistics; ii) generation of a ~1.56 kb RT-PCR or PCR amplicon; and, iii) bi-directional sequencing of the amplicon by the Sanger method (4, 5).

**NEAR-FULL LENGTH (NFL) SINGLE PROVIRAL AND VIRAL GENOME AMPLIFICATION AND SEQUENCING**

Near-full length (NFL) HIV-1 DNA was amplified by nested PCR from genomic DNA extracted from PBMC or total $CD4^+$ T-cell at a proviral endpoint of a single template per PCR reaction as determined by Poisson distribution statistics. PCR amplifications were performed using the 2x RANGER DNA Polymerase Mix according to the manufacturer's recommendations (Bioline) and previously reported primers (6). The sizes of the NFL amplicons were confirmed with the Perkin Elmer GX Touch 24 LabChip bioanalyzer using the 12K DNA module. NFL amplicons containing 16bp symmetrical barcodes were size selected by BluePippin (Sage Science) and libraries were constructed using the PacBio SMRTbell Template Prep Kit prior to PacBio sequencing (Pacific Biosciences). Alternatively, NFL amplicons were sequenced by Illumina MiSeq using the KAPA HyperPlus kit (KAPA Biosystems) for library construction and the MiSeq nano v2 500 cycle, 2x250run kit with dual index (Illumina) for sequencing. SGS of overlapping half genomes from virion-associated HIV-1 RNA in p24$^+$ qVOA wells was performed as reported (4, 7).

**ASSESSMENT OF CELL CLONALITY**

A clone of HIV-1 infected cells was defined by identifying multiple cells that had identical proviral sequences integrated into the identical position in the human genome. The methods used to confirm clonality are described below.

**MULTIPLE-DISPLACEMENT AMPLIFICATIONS (MDA) OF GENOMIC DNA**

Whole cellular genome amplification was done as previously described (8) with the following modifications: 1) genomic DNA was diluted to a proviral endpoint of one template per reaction; ii) reactions volumes were 25μl containing D solution [final concentrations: 24.4mM KOH and 20μM random hexamers (OH-5'-NNNN*N*N-3')], N solution [final concentrations: 25mMTrizma-HCl, pH 7.5 and 600mM Trehalose], and reaction mix [final concentrations: 1 X Phi buffer, 20mM KCl, 2mM DTT, 100mM BSA, 1.8mM dNTPs, 600mM Trehalose, and 1unit phi29 DNA polymerase]; and, iii) incubations were performed at 40$^0$C for 20h and terminated at 65$^0$ C for 10 minutes.

**INTEGRATION SITE ANALYSIS**

Integration site analyses (ISA) were conducted as previously described (9). For integration site analysis of proviruses with specific viral sequences of interest (i.e., matching plasma HIV-1 RNA sequences), ISA was performed with the following modifications: The starting template was 0.8× SPRI purified MDA material that contained the *gag-pro-pol* sequence that matched the plasma HIV-1 RNA or p24$^+$ qVOA HIV-1 RNA sequence of interest. ISA was performed using an in-house workflow utilizing multiple displacement amplification and a specificity-enhancing linker-mediated PCR that amplifies across the 5'LTR host/virus junction (10).

**HOST-FULL LENGTH PROVIRUS-HOST AMPLIFICATION AND SEQUENCING (HFH)**

For specific clones of interest, full-length sequences (host to full-length provirus to host [HFH]) of the proviruses integrated at the same site were assessed for identity. Proviral integration site positions are based on data in the Human Genome Browser at UCSC using the human genome assembly reference Hg19 sequence (11, 12). These sequences were also used to design primers for the human host sequences adjacent to full length proviruses. Host-specific primers were used with SGS *gag-pro-pol* proviral specific primers to amplify the entire provirus and the flanking 5' and 3' host sequences as two overlapping fragments from genomic DNA extracted from PBMC or total $CD4^+$ T-cells using primers specific for each integrant of interest (Table S4). The amplification was performed using 2x RANGER DNA Polymerase Mix (Bioline) as described above for NFL proviral amplification. The products were sequenced using either Sanger or the Illumina platform. Primers used for Sanger sequencing are listed in Table S5.

**SEQUENCE ALIGNMENTS, QUALITY CONTROL, AND PHYLOGENETIC ANALYSES**

Sequence alignments and phylogenetic analyses of the *gag-pro-pol* sequences were performed as reported previously (4). This included alignments, exclusion of mixtures, and quality control of sequences in Sequencher v5.0 (Gene Codes) (Figure S6). Neighbor-joining *p*-distance phylogenetic trees were rooted to subtype B with bootstrapping at 1,000 replicates per tree using MEGA6 (13).

**HIV-1 SUBTYPE, DRUG SUSCEPTIBILITY ANALYSES, DRUG CONCENTRATION DETERMINATION AND CO-RECEPTOR TROPISM**

HIV-1 subtype and genotype-predicted susceptibilities to antiretroviral drugs were determined by the HIVdb algorithm (Stanford University HIV Drug Resistance Database) (14). Drug level concentrations in human plasma for the non-nucleoside reverse transcriptase inhibitor efavirenz and the protease inhibitors darunavir, atazanavir, and ritonavir, were measured by a gradient separation through Ultra Performance Liquid Chromatography with Electrospray Ionization Tandem Mass Spectrometry for detection (15). Drug level concentration in human plasma for the integrase inhibitor dolutegravir was measured using a protein precipitation method with isocratic separation by Liquid Chromatography with tandem mass spectrometry for detection (16). Co-receptor tropism was determined by the Geno2Pheno bioinformatics software using both established and individually selected cutoffs (17).

**IMMUNOPHENOTYPING**

Surface flow cytometric staining of CD3, CD4, CD8, CD19, CD56, CD25, CD69, CD38, HLA-DR, and CD107a (BD Biosciences) was performed on a BD LSRII cytometer according to standard published methods (18), analyzed using FlowJo, and results compared to published results of healthy adults (19-26).

**STATISTICAL ANALYSES**

Average pairwise distances were calculated by MEGA6 through the exclusion of hypermutants and identification of hypermutants was performed using the HIVdb algorithm (Stanford University HIV Drug Resistance Database) (14).

**DATA AVAILABILITY**

Sequences are being submitted to the GenBank database. Viral sequences and integration sites were also submitted to the Proviral Sequence Database (PSD) and the Retrovirus Integration Database (RID;

https://rid.ncifcrf.gov/) at the U.S. Department of Health and Human Services, National Institutes of Health, National Cancer Institute (27).

#### Figure S1: Panels A-F

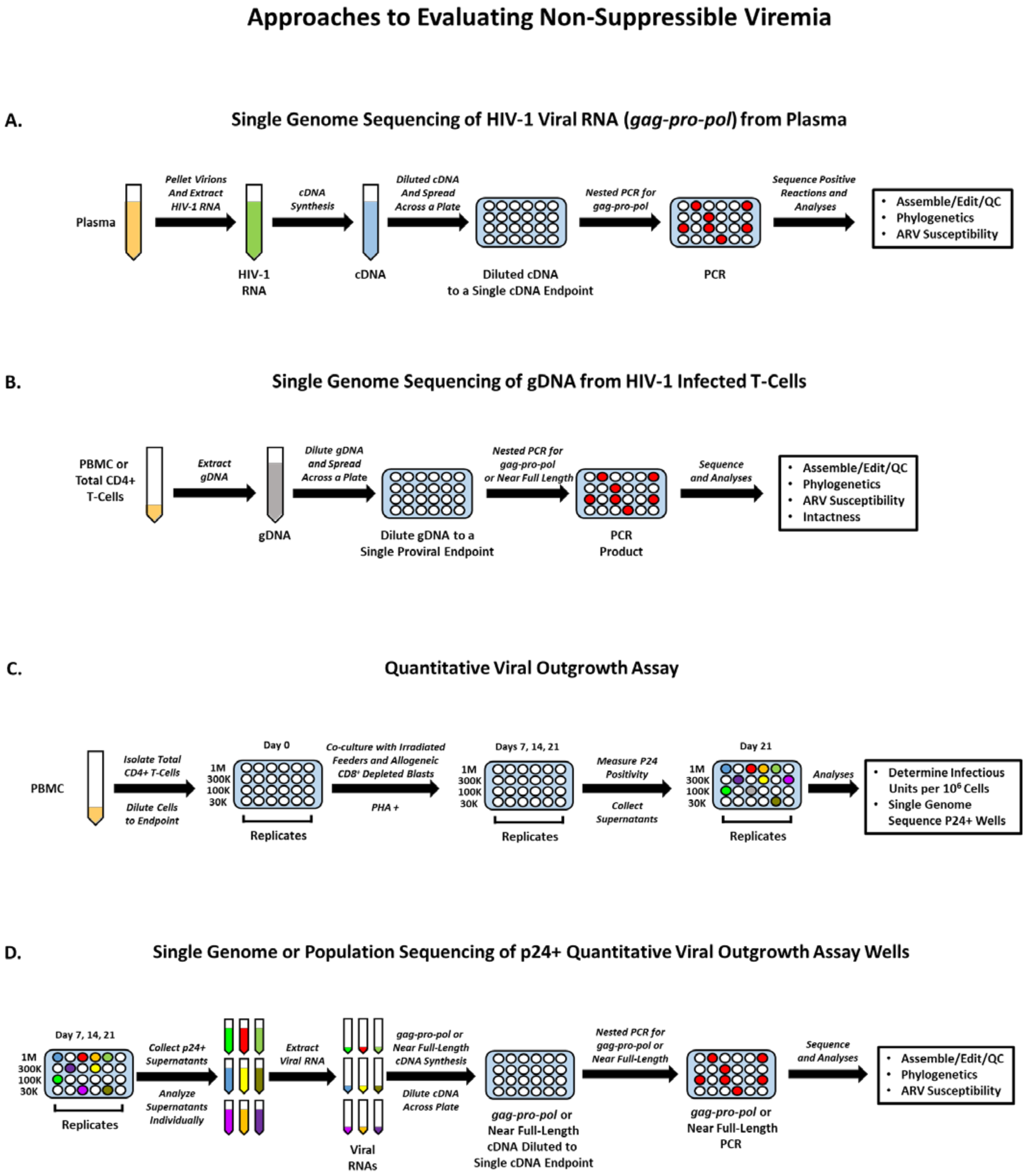

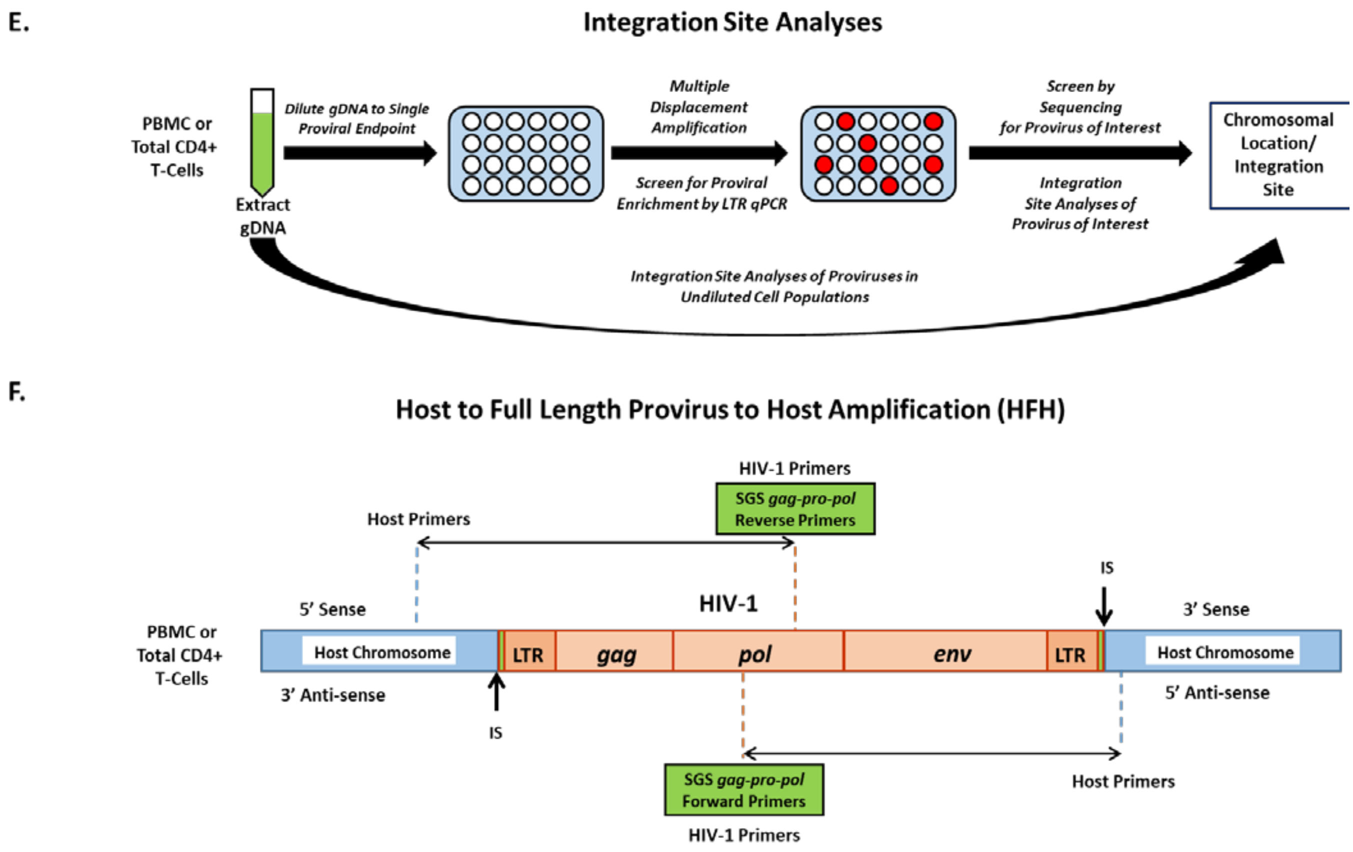

**Figure S1. Schematic of Experimental Approaches to Evaluate Non-Suppressible Viremia.** Donor plasma and PBMC or total CD4$^+$ T cells originated from large volume blood draws or leukaphereses. Single genome sequencing (SGS) of HIV-1 RNA from plasma generated an amplicon containing a portion of *gag (p6)*, all of *pro*, and, from *pol* the portion encoding the 1$^{st}$ 300 amino acids of reverse transcriptase (*gag-pro-pol*) (Panel A). Proviral HIV-1 DNA SGS of *gag-pro-pol* or near-full length amplicons from PBMC or total CD4$^+$ T cells (Panel B). Quantitative viral outgrowth assays performed using total CD4$^+$ T cells (Panel C). SGS of *gag-pro-pol* or near-full length HIV-1 RNA genome from quantitative viral outgrowth assay p24$^+$ wells (Panel D). Integration site analyses (ISA) were preformed directly on undiluted cell populations or at a single proviral endpoint using multiple displacement amplification of DNA originating from PBMC or total CD4$^+$ T cells (Panel E) (9). For proviruses of interest, host-to-full length provirus-to host amplification and sequencing was used to confirm the sequence identity of the clonally-expanded provirus (Panel F).

#### Figure S2

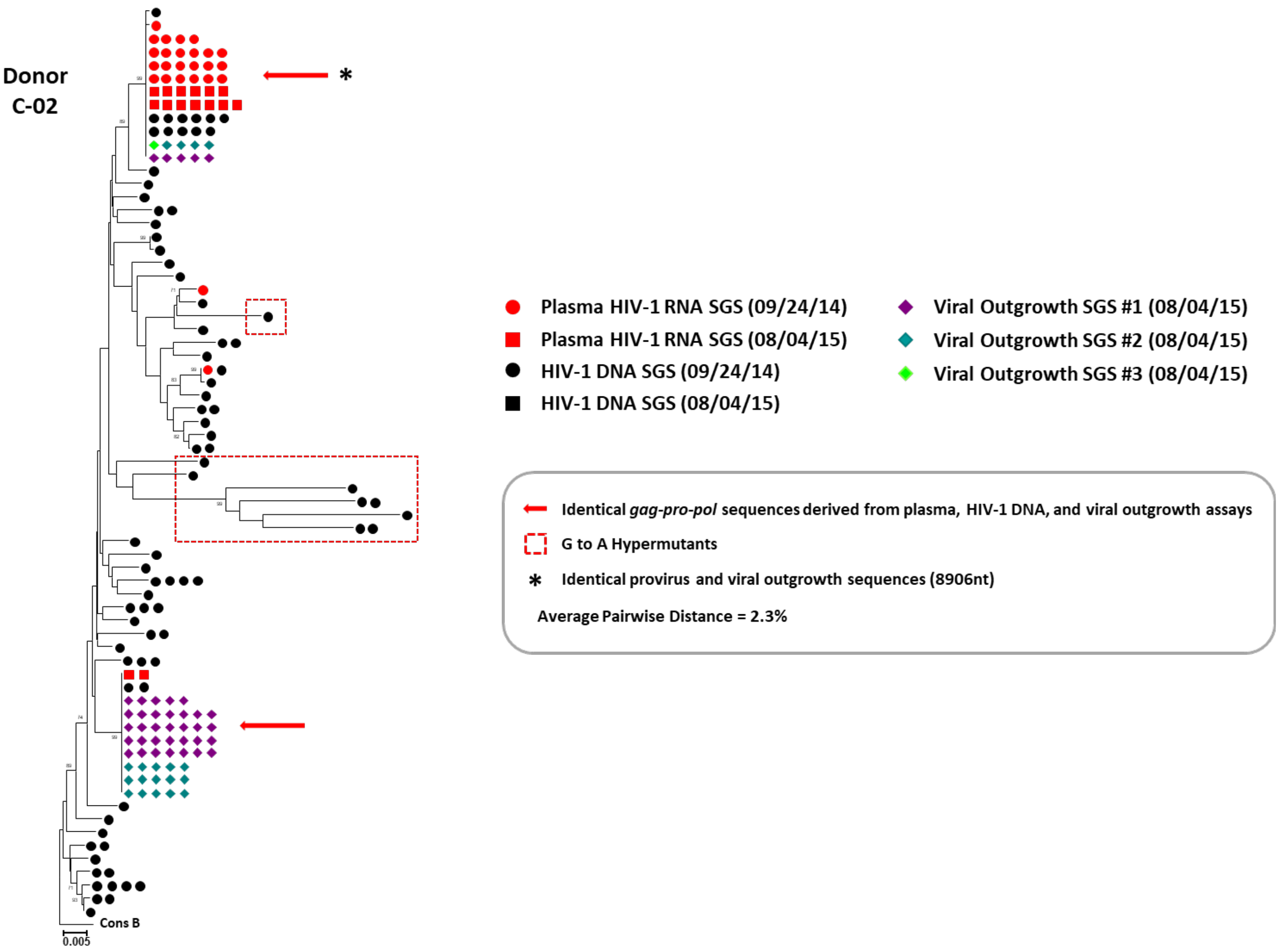

**Figure S2. Representative Neighbor-Joining *p*-Distance Phylogenetic Tree of Plasma HIV-1 RNA-, HIV-1 DNA-, and Quantitative Viral Outgrowth Assay (qVOA)-Derived Sequences from Donor C-02 with Intact Replication-Competent Proviruses Producing Non-Suppressible Viremia.** The trees were rooted to a subtype B consensus sequence. Single genome sequences (SGS) of a portion of *gag (p6)*, all of *pro*, and the portion of *pol* that encodes the 1st 300 amino acids of reverse transcriptase (*gag-pro-pol*) (5) were obtained from plasma HIV-1 RNA, HIV-1 DNA from peripheral blood mononuclear cells, and culture supernatants from p24+ qVOA wells for donor C-02. Red circles and squares represent plasma-derived sequences from two different time points. Black circles and squares represent HIV-1 DNA-derived sequences from two different time points. Different colored diamonds represent viral outgrowth assay-derived sequences from independent p24$^+$ wells. A red arrow shows identical *gag-pro-pol* sequences for plasma-, HIV-1 DNA-, and viral outgrowth assay HIV-1 RNA-derived sequences. The asterisk shows matching sequences of provirus (near-full length HIV-1 DNA and host to full length provirus to host amplicons) and near-full length viral RNA sequences from p24$^+$ wells. HIV-1 DNA sequences with G to A hypermutations are enclosed in red-hashed boxes. The viral outgrowth sequence variants that differ by 1-2 nucleotides can be attributed to either ex vivo replication or errors introduced during cDNA synthesis. Average pairwise distances (APD) calculated by MEGA v6.0 using HIV-1 DNA sequences and excluding hyper-mutated sequences.

#### Figure S3

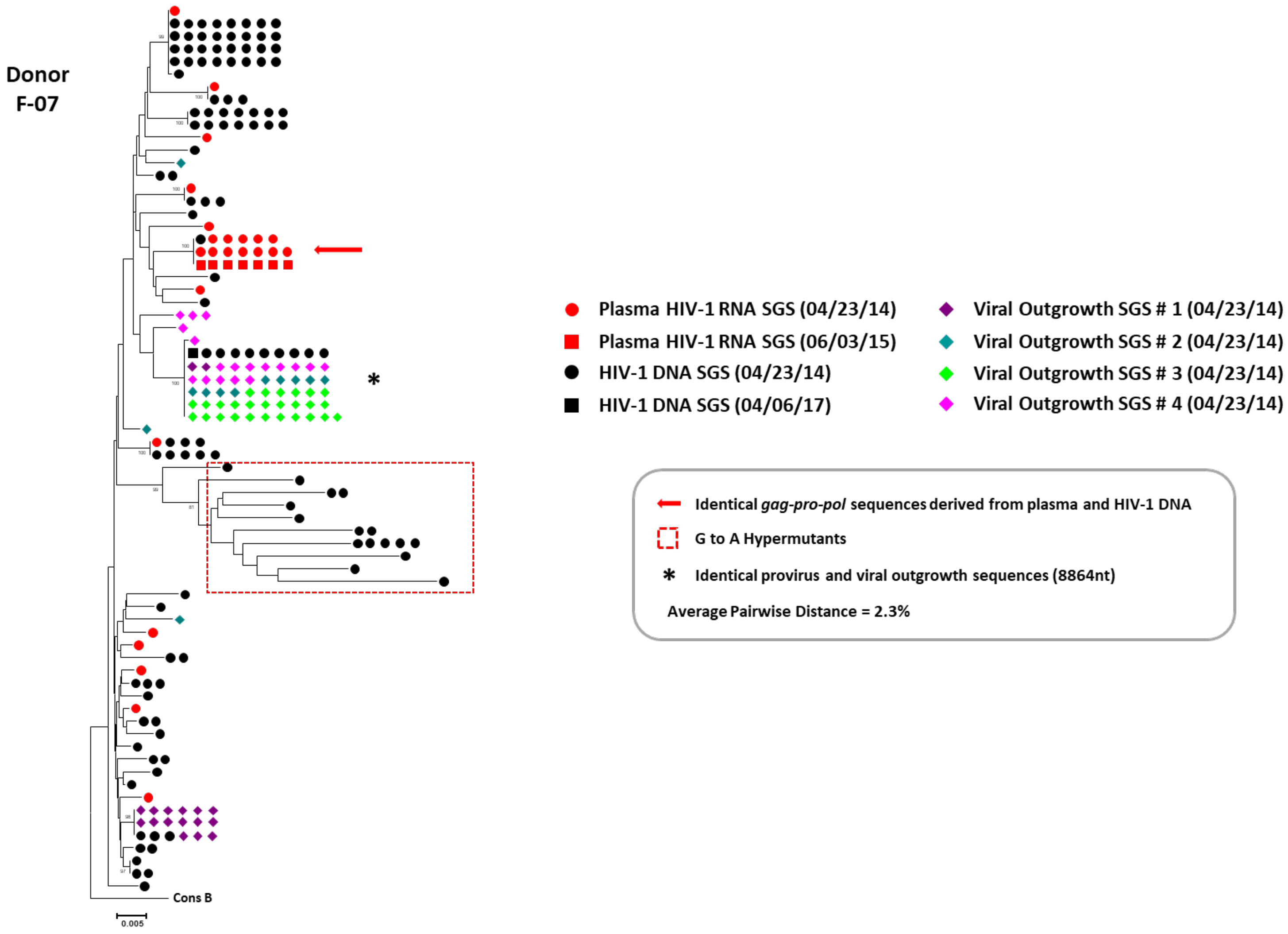

**Figure S3. Representative Neighbor-Joining *p*-Distance Phylogenetic Tree of Plasma HIV-1 RNA-, HIV-1 DNA-, and Quantitative Viral Outgrowth Assay (qVOA)-Derived Sequences for Donor F-07 with an Intact Replication Competent Provirus.** The trees were rooted to a subtype B consensus sequence. Single genome sequences (SGS) of a portion of *gag (p6)*, all of *pro*, and the portion of *pol* encoding the 1$^{st}$ 300 amino acids of reverse transcriptase (*gag-pro-pol*) (5) were obtained from plasma HIV-1 RNA, HIV-1 DNA from peripheral blood mononuclear cells (PBMC), and culture supernatants from p24$^{+}$ qVOA wells for donor F-07. Red circles and squares represent plasma-derived sequences from two different time points. Black circles and squares represent HIV-1 DNA-derived sequences from two different time points. Different colored diamonds represent viral outgrowth assay-derived sequences from independent p24$^+$ wells. A red arrow shows identical *gag-pro-pol* sequences from plasma-, HIV-1 DNA-, and viral outgrowth assay HIV-1 RNA-derived sequences. The asterisk shows matching sequences of provirus (near-full length HIV-1 DNA and host through full length provirus to host amplicons) and near-full length viral RNA sequences from p24$^+$ wells. HIV-1 DNA sequences with G to A hypermutations are enclosed in red-hashed boxes. The viral outgrowth sequence variants that differ by 1-2 nucleotides can be attributed to either ex vivo replication or errors introduced during cDNA synthesis. Average pairwise distances (APD) calculated by MEGA v6.0 using HIV-1 DNA sequences and excluding hyper-mutated sequences.

#### Figure S4: Panels A-D

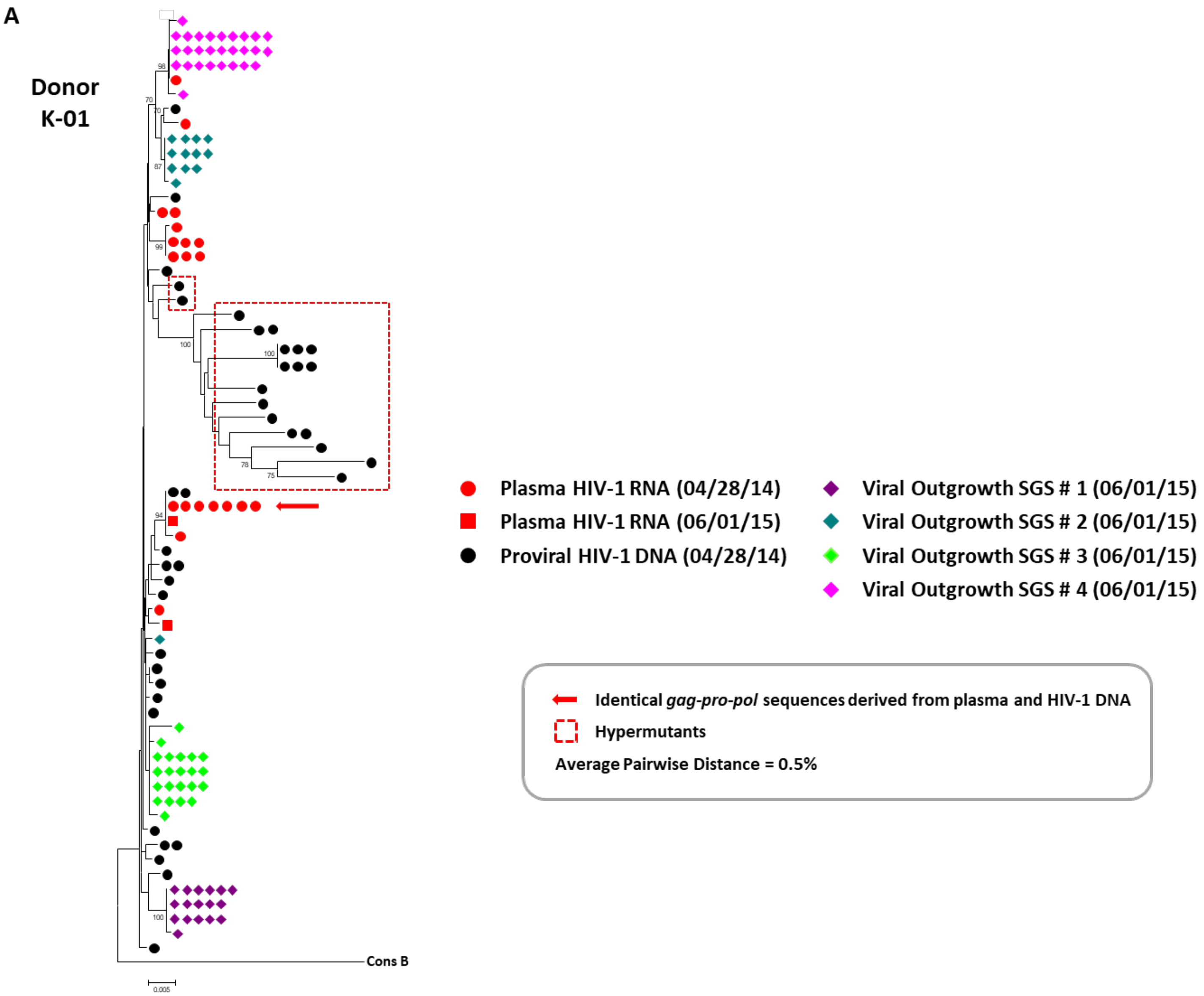

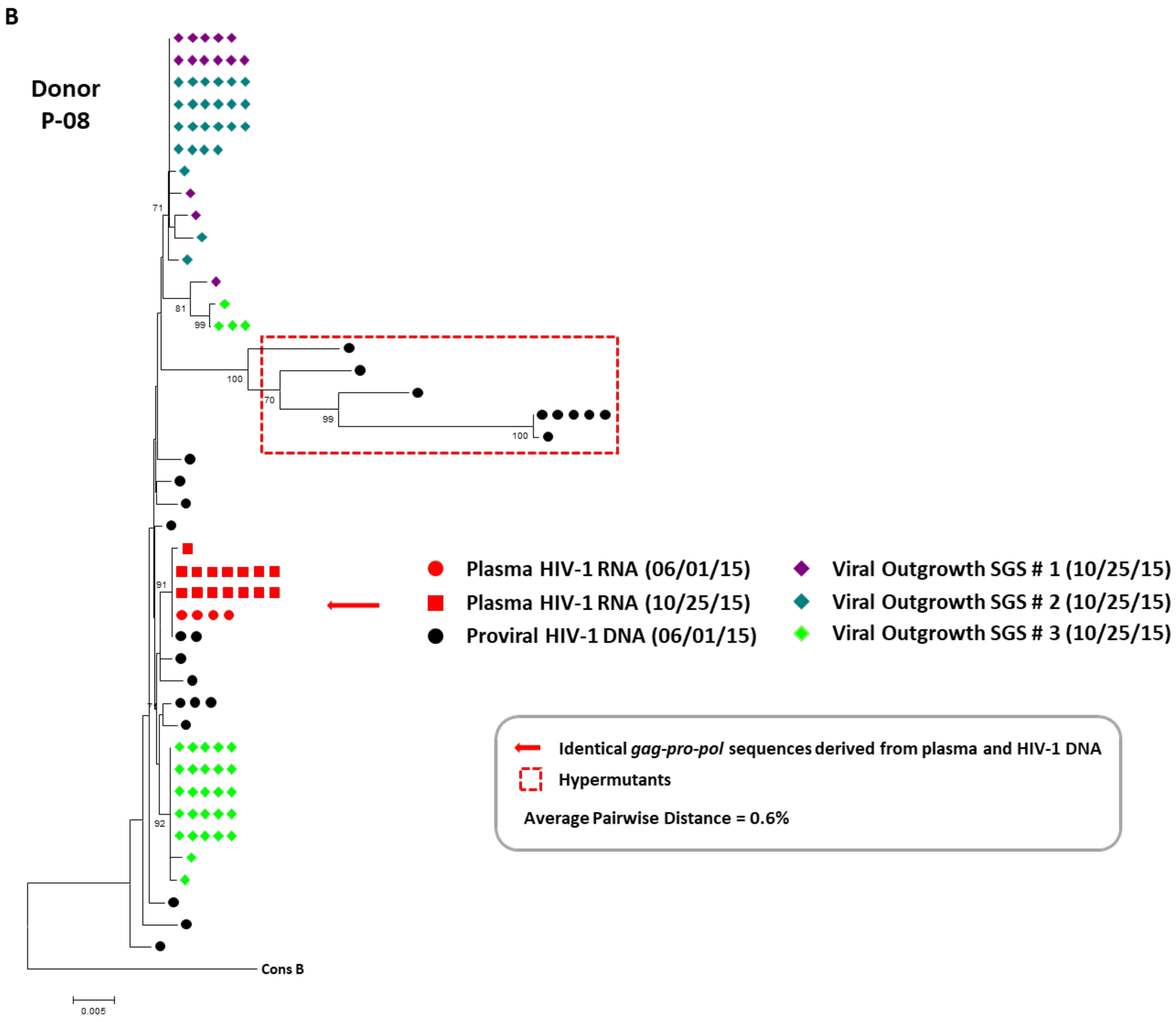
B
Donor
P-08
71
81
99
100
70
99
100
91
92
Cons B
0.005
Plasma HIV-1 RNA (06/01/15)
Plasma HIV-1 RNA (10/25/15)
Proviral HIV-1 DNA (06/01/15)
Viral Outgrowth SGS # 1 (10/25/15)
Viral Outgrowth SGS # 2 (10/25/15)
Viral Outgrowth SGS # 3 (10/25/15)
Identical gag-pro-pol sequences derived from plasma and HIV-1 DNA
Hypermutants
Average Pairwise Distance = 0.6%

C

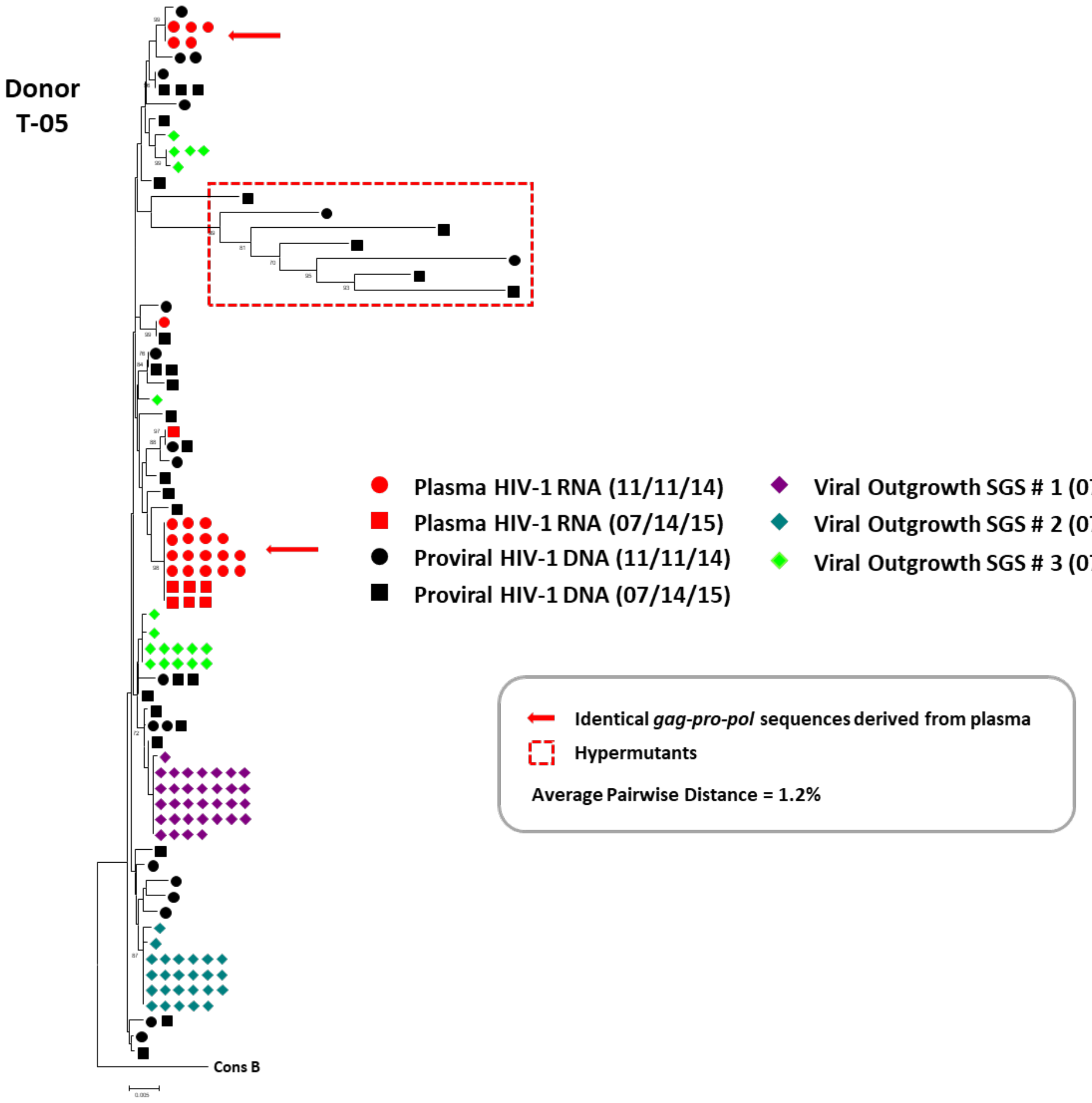
Donor
T-05
Cons B
Plasma HIV-1 RNA (11/11/14)
Plasma HIV-1 RNA (07/14/15)
Proviral HIV-1 DNA (11/11/14)
Proviral HIV-1 DNA (07/14/15)
Viral Outgrowth SGS # 1 (07/14/15)
Viral Outgrowth SGS # 2 (07/14/15)
Viral Outgrowth SGS # 3 (07/14/15)
Identical gag-pro-pol sequences derived from plasma
Hypermutants
Average Pairwise Distance = 1.2%

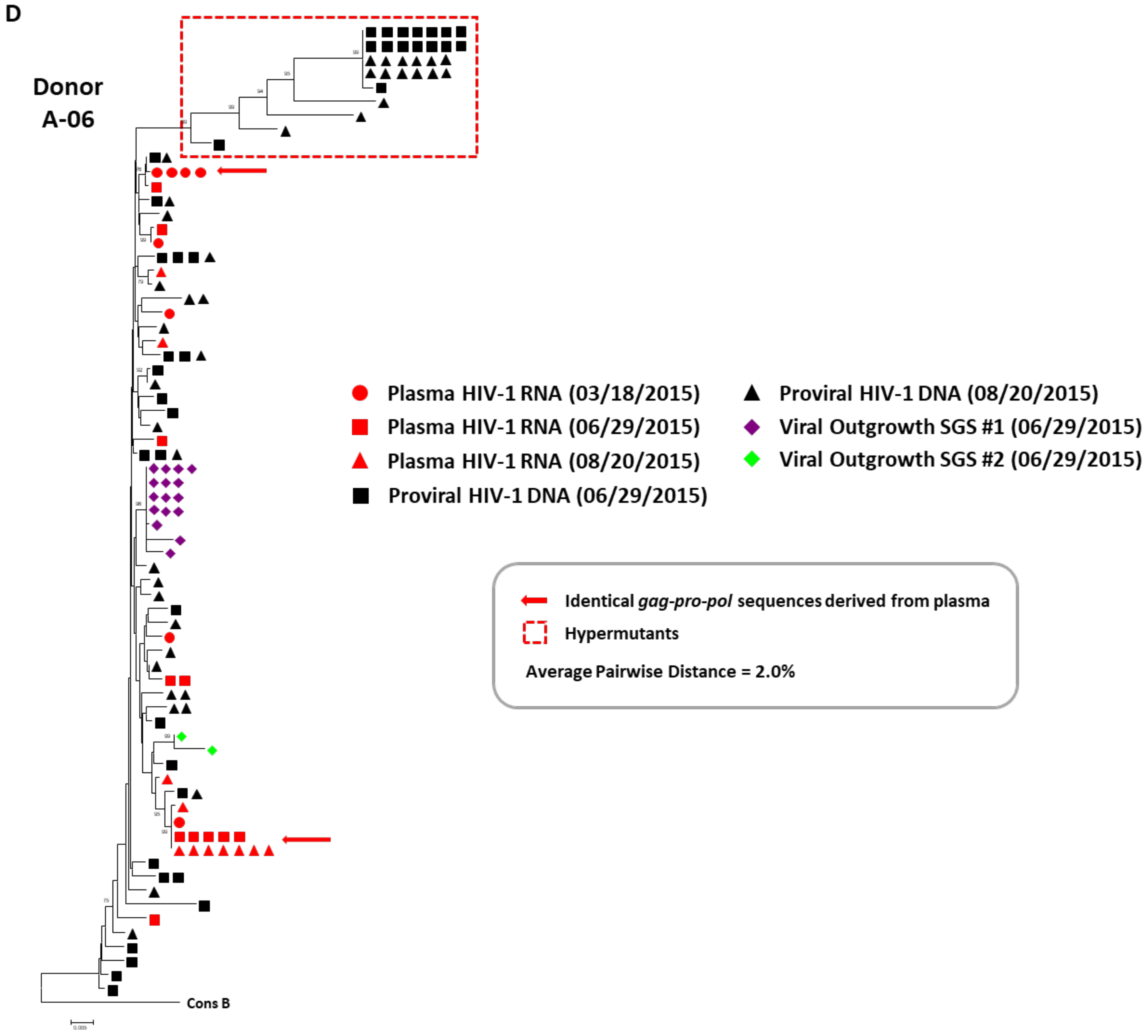

**Figure S4. Representative Neighbor-Joining *p*-Distance Phylogenetic Trees of Plasma HIV-1 RNA-, HIV-1 DNA-, and Quantitative Viral Outgrowth Assay (qVOA)-Derived Sequences for Donors K-01, P-08, T-05, and A-06 with Non-Suppressible Viremia.** The trees were rooted to a subtype B consensus sequence. Single genome sequences (SGS) of a portion of *gag (p6)*, all of *pro*, and the portion of *pol* encoding the 1st 300 amino acids of reverse transcriptase (*gag*-

*pro-pol*) (5) were obtained from plasma HIV-1 RNA, HIV-1 DNA from peripheral blood mononuclear cells (PBMC), and culture supernatants from p24$^+$ qVOA wells for donors K-01 (Panel A), P-08 (Panel B), T-05 (Panel C), and A-06 (Panel D). Red circles, squares, and triangles represent plasma-derived sequences from three different time points. Black circles and squares represent HIV-1 DNA-derived sequences from two different time point. Different colored diamonds represent viral outgrowth assay-derived sequences from independent p24$^+$ wells. A red arrow shows identical *gag-pro-po*l sequences for plasma HIV-1 RNA- and HIV-1 DNA-derived sequences for donors K-01 and P-08 or only plasma HIV-1 RNA-derived sequences for donors T-05 and A-06. HIV-1 DNA sequences with G to A hypermutations are enclosed in red-hashed boxes. The viral outgrowth sequence variants that differ by 1-2 nucleotides can be attributed to either ex vivo replication or errors introduced during cDNA synthesis. Average pairwise distances (APD) calculated by MEGA v6.0 using HIV-1 DNA sequences and excluding hyper-mutated sequences.

#### Figure S5

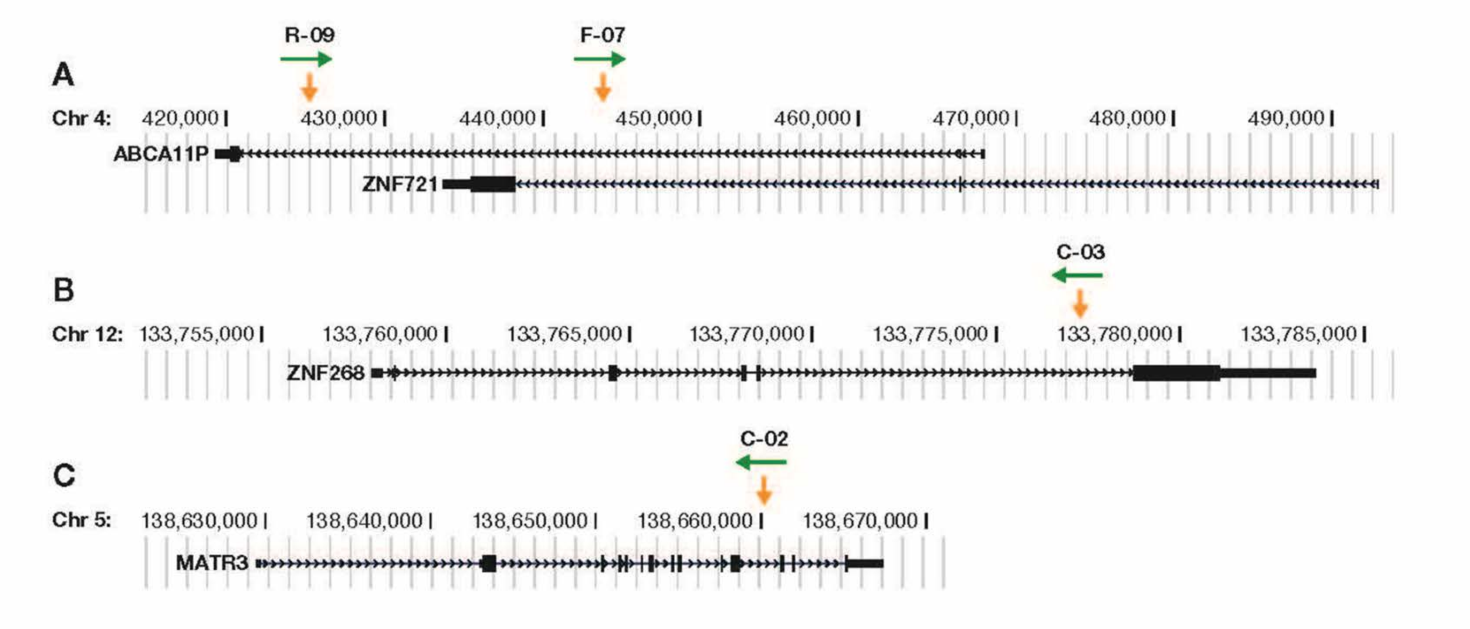

**Figure S5. Gene Diagrams with Integrated, Intact, Infectious Proviruses.** The gene diagrams are based on the hg19 reference of the human genome and were prepared using the UCSC genome browser (https://genome.ucsc.edu/). There are, for each of the four genes shown in the diagram, additional transcripts that are not shown. The small black arrowheads along the black lines in the gene diagrams denote the direction in which the genes are transcribed. Exons are shown as bars within the gene. The narrow portions of the exons at the ends of the genes are non-translated regions; the wider exons/portions of exons are coding. The yellow vertical arrows mark the sites where the proviruses were integrated; the horizontal green arrows show the direction in which the proviruses were oriented. The organization of the overlapping genes *ABCA11P* and *ZNF721* and the sites of integration for the repliclone proviruses from donors R-09 and F-07 (Panel A). The organization of the *ZNF268* gene and integration site of the repliclone provirus from donor C-03 (Panel B). The organization of the *MATR3* gene and integration site of the repliclone provirus from donor C-02 (Panel C).

**Figure S6**

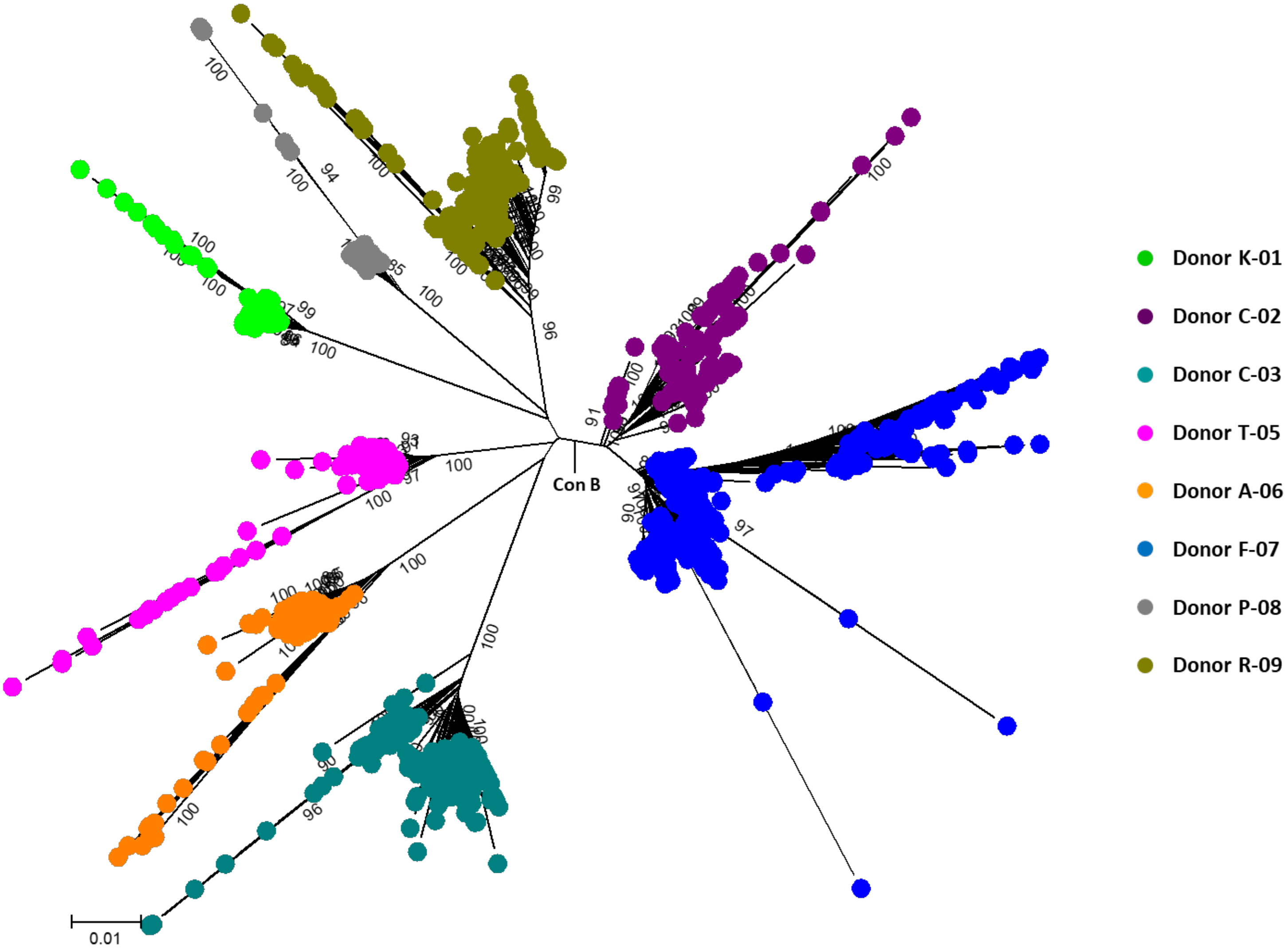

**Figure S6. Neighbor-Joining *p*-Distance Radial Tree of All Sequences from All Donors.** The tree was rooted to a subtype B consensus sequence. Single genome sequences (SGS) of a portion of *gag (p6)*, all of *pro*, and the portion of *pol* encoding the 1st 300 amino acids of reverse transcriptase (*gag-pro-pol*) (5) were obtained from plasma HIV-1 RNA, HIV-1 DNA in peripheral blood mononuclear cells (PBMC), and culture supernatants from p24+ qVOA wells. All sequences from each donor are represented as different colored circles; Donor K-01 (Green), C-02 (Purple), C-03 (Teal), T-05 (Pink), A-06 (Orange), F-07 (Blue), P-08 (Grey), and R-09 (Gold).

**Table S1.** HIV Drug Susceptibilities, Diversity, and Coreceptor Tropisms

| Donor ID | Mutations Causing Resistance to Donor's Current ART Regimen[a] | Average Pairwise Distance of All Proviruses(%)[b] | Geno2pheno Coreceptor Tropisms[c,d] |
|---|---|---|---|
| R-09 | 0/30 | 2.4 | R5-Tropic (13.2%FPR) |
| C-03 | 0/49, 0/13* | 1.9 | R5-Tropic (62.8%FPR) |
| C-02 | 0/47 | 2.3 | R5-Tropic (48.7%FPR) |
| F-07 | 0/39[†] | 2.3 | R5-Tropic (33%FPR) |
| K-01 | 0/24 | 0.5 | - |
| P-08 | 0/21 | 0.6 | - |
| T-05 | 0/34 | 1.2 | - |
| A-06 | 0/31 | 2.0 | - |

[a] Drug-resistance mutations in plasma were identified using the Stanford HIV Drug Resistance Database v8.7; genotype covered protease, reverse transcriptase and integrase in individuals receiving an integrase inhibitor*

[b] Average pairwise distance of all proviral sequences calculated using MEGA v6.0

[c] Coreceptor tropisms predicted by Geno2pheno for X4/R5 tropism using near-full length sequences

[d] R5, CCR5-tropic; X4, CXCR4-tropic; FPR, false-positive rate for incorrectly identifying the sequence as X4-tropic was determined using cutoffs either from the German (5% to 15% cutoff), European (10% cutoff) and MOTIVATE (2% to 5.75% cutoff) guidelines. False positive rate (FPR) results represented are using European Guideline cutoffs of which were identical to those obtained by German and MOTIVATE guidelines

[†] 21 of 39 plasma-derived sequences contained D67N/K70R/K219Q conferring low-level resistance by the Stanford HIVdb algorithm to Abacavir

**Table S2.** Antiretroviral Drug Levels in Individuals with Non-suppressible Viremia

| Donor | Sample Type | Sample Draw Date | Patient Regimen[a] | DRV (ng/mL) | RTV (ng/mL) | ATV (ng/mL) | EFV (ng/mL) | DTG (ng/mL) |
|---|---|---|---|---|---|---|---|---|
| R-09 | Plasma | 6/4/2015 | TDF/FTC/**EFV** | - | - | - | 1636 | - |
| R-09 | Plasma | 8/6/2015 | TDF/FTC/**EFV** | - | - | - | 1334 | - |
| C-03 | Plasma | 9/30/2014 | **DRV/r**/ETV/**DLG** | 4522 | 441 | - | - | 2033 |
| C-02 | Plasma | 9/24/2014 | TDF/FTC/**EFV** | - | - | - | 3321 | - |
| F-07 | Plasma | 6/3/2015 | ABC/3TC/**EFV** | - | - | - | 6766 | - |
| K-01 | Plasma | 4/28/2014 | TDF/FTC/**DRV/r** | 2277 | 115 | - | - | - |
| P-08 | Plasma | 9/29/2015 | TDF/FTC/**EFV** | - | - | - | 975 | - |
| T-05 | Plasma | 11/11/2014 | TDF/FTC/**ATV/r** | - | 36.5 | 655 | - | - |
| A-06 | Plasma | 1/29/2015 | TDF/FTC/**ATV/r** | - | 177 | 1636 | - | - |
| | | | **Target Trough Range ng/mL** | 1000-8000 | <50-2500 | 150-850 | 1000-4000 | 800-1000 |

[a] Antiretroviral drug abbreviations: ABC (abacavir); ATV/r (atazanavir/ritonavir; DRV/c (darunavir/cobicistat); DRV/r (darunavir/ritonavir; DTG (dolutegravir); EFV (efavirenz); ETV (etravirine); 3TC (lamivudine); FTC (emtricitabine); TDF (tenofovir disoproxil fumarate)
Antiretroviral drugs tested are shown in **Bold**
DRV, EFV assay range (200-15,000 ng/mL); ATV, RTV assay range (10-4,000 ng/mL); DTG assay range (20-10,000 ng/mL)

**Table S3.** Immunophenotyping of PBMC from Donors Referred for Non-Suppressible Viremia

| | **Frequency (%) of Lymphocytes** | | | | | | **Cellular Activation Markers** | | | | |
|---|---|---|---|---|---|---|---|---|---|---|---|
| | | | | | | | **Frequency (%) of CD4$^+$ T-Cells** | | | | **Frequency (%) of CD8$^+$ T-Cells** |
| ***Donor ID*** | ***CD3$^+$ CD4$^+$*** | ***CD3$^+$ CD8$^+$*** | ***CD3$^+$ CD4$^+$ CD8$^+$*** | ***CD3$^-$ CD19$^+$*** | ***CD3$^-$ CD56$^+$*** | | ***CD25$^+$*** | ***CD69$^+$*** | ***HLA-DR$^+$*** | ***CD38$^+$ HLA-DR$^+$*** | ***CD107a$^+$*** |
| R-09 | 31.8 | 12.5 | 0.87 | 27.8 | 4.7 | | 9.8 | 12.4 | 22.2 | 7.9 | 2.6 |
| C-03 | 31.7 | 31.7 | 0.72 | 15.8 | 2.1 | | 9.3 | 6.4 | 9.0 | 4.2 | 0.7 |
| C-02 | 33.5 | 26.4 | 0.42 | 6.5 | 8.5 | | 8.7 | 19.6 | 16.4 | 8.0 | 3.9 |
| F-07 | 26.1 | 33.0 | 1.08 | 8.1 | 1.9 | | 5.2 | 3.2 | 10.3 | 5.3 | 0.8 |
| K-01 | 30.9 | 29.3 | 1.05 | 10.4 | 5.2 | | 13.1 | 3.2 | 18.4 | 5.4 | 0.4 |
| P-08 | 17.0 | 37.7 | 0.75 | 9.4 | 11.2 | | 6.5 | 9.4 | 31.7 | 8.7 | 2.6 |
| T-05 | 48.5 | 22.9 | 1.26 | 5.9 | 4.1 | | 8.7 | 1.5 | 5.9 | 3.0 | 1.0 |
| A-06 | 32.0 | 32.0 | 0.69 | 5.9 | 1.1 | | 10.5 | 4.1 | 19.7 | 4.1 | 1.5 |
| **Median** | 31.8 | 30.5 | 0.75 | 8.8 | 4.4 | | 9.0 | 5.3 | 17.4 | 5.4 | 1.3 |
| **Healthy Donor** | 32.5-68.3[a] | 11.5-38.6[a] | 0.25-6[b] | 4.7-22.5[a] | 6.3-10[c] | | 0.3-10.7[a] | 1-9[d] | 0.8-4.4[a] | 0.3-1.35[a] | 2-20[e] |

[a] Range (%) of lymphocyte subsets CD3$^+$/CD4$^+$, CD3$^+$/CD8$^+$, CD3$^-$/CD19$^+$, CD4$^+$/CD25$^+$, CD4$^+$/HLA-DR$^+$, and CD4$^+$/CD38$^+$/HLA-DR$^+$ in healthy human adults (19,20)

[b] Range (%) of lymphocyte subset CD3$^+$/CD4$^+$/CD8$^+$ in healthy human adults (21,22)

[c] Range (%) of lymphocyte subset CD3$^-$/CD56$^+$ in healthy human adults (23,24)

[d] Range (%) of lymphocyte subset CD4$^+$/CD69$^+$ in healthy human adults (25)

[e] Range (%) of lymphocyte subset CD8$^+$/CD107a$^+$ in healthy human adults (26)

**Table S4.** Host to Full Length Provirus to Host Amplification Primer Sets

| Donor (Provirus) | HIV-Specific Primers | HIV-Specific Primer Sequences (5'-3') | Host-Specific Primer | Host-Specific Primer Sequences (5'-3') |
|---|---|---|---|---|
| R-09 (ABCA1P) | RF-09_F1 | GATGACAGCATGCCAGGGAG | ABCA11P_R1 | TGGGATTACAGGCTGGGATAATG |
| | RF-09_F2 | GAGTCTTAGCTGAAGCAATGAG | ABCA11P_R2 | GTATAACGTAAAATGAATACATCCTTGTC |
| | RF-09_R1 | CTATTAAGTCTTTTGATGGGTCATAG | ABCA11P_F1 | TGGTTGTTCCCTATACATTTTAATC |
| | RF-09_R2 | CTGTTAGTGGTATTACTTCTGTTAGTGCTT | ABCA11P_F2 | ATTCTCAGTGTAGAGCGTGGTTACC |
| C-03 (ZNF268) | CN-03_F1 | GATGACCGCATGTCAGGGAG | ZNF268_F1 | CACAAAGCTGTTTGCCTACCC |
| | CN-03_F2 | GAGTCTTGGCTGAAGCAATGAG | ZNF268_F2 | TTCTTTTCCATGCCTGCTAGAG |
| | CN-03_R1 | CTATTAAATCTTTTGATGGGTCATAA | ZNF268_R1 | GCAGAGAACAATGCAGATTACT |
| | CN-03_R2 | CTGTCAGTGGTACTACATCTGTTAGTGCTT | ZNF268_R2 | CAGGATAAAAATTGCACAGCAGGC |
| C-02 (MATR3) | CF-02_F1 | GATGACCGCATGTCAGGGAG | MATR3_F1 | CCCAACATAGTTAAAACTTTTGCCACTCATTC |
| | CF-02_F2 | GAGTCTTGGCTGAAGCAATGAG | MATR3_F2 | ATTTGAACAAGTAAGTCATTTAGAAGCC |
| | CF-02_R1 | CTATTAAATCTTTTGATGGGTCATAA | MATR3_R1 | GGAAACGGATAGCGTCTTTG |
| | CF-02_R2 | CTGTCAGTGGTACTACATCTGTTAGTGCTT | MATR3_R2 | GGAAGGTCTGCCTCACACAAAG |
| F-07 (ZNF721/ ABCA11P) | FN-07_F1 | GATGACAGCATGTCAGGGAG | ZNF721_R1 | CTCTTAAAGTCTCTTGCCTATATTCAAATTG |
| | FN-07_F2 | GGGTTTTGGCGGAAGCAATGAG | ZNF721_R2 | CTGCCTGTGCTTTTGAGGTCTTAAG |
| | FN-07_R1 | CTATTAAGTCTTTTGATGGGTCATAA | ZNF721_F1 | GGTGCTAGGAAAATTATCTACAAG |
| | FN-07_R2 | TTGTTAGTGGTACTACTTCTGTTAGTGCCT | ZNF721_F2 | GAAACCCAAATAAAGCTATTTGAAGTAAACATAC |

**Table S5.** Sequencing Primers Used for Near-full Proviral Sequencing

| Sequencing Primer Name | Primer Sequence | Sequencing Primer Name | Primer Sequence |
|---|---|---|---|
| **A** | 5'-CTTTCGCTTTCAAGTCCCT-3' | **M** | 5'-AAGAGATATAGCACACAAGTAG-3' |
| **B** | 5'-AAATCTCTAGCAGTGGCG-3' | **M_R-09** | 5'-AGGAACTATAGCACACAAGTAG-3' |
| **SGS_C** | 5'-TTCTTCTGTCAATGGCCATTGTTTAAC-3' | **N** | 5'-ATTGGGTGCCAACATAGCAGAATA-3' |
| **SGS_D** | 5'-TTGCCCAATTCAATTTTCCCACTAA-3' | **O** | 5'-CTATGGCAGGAAGAAGCGG-3' |
| **C** | 5'-CAAAGGATAGAGGTAAAAGAC-3' | **P** | 5'-GTGGAAAAATAACATGGTAG-3' |
| **C_R-09** | 5'-CAAAGGATAGATGTAAAAGAT-3' | **Q** | 5'-CAGTAGTATCAACTCAACTG-3' |
| **C_R-09_RC** | 5'-ATCTTTTACATCTATCCTTTG-3' | **Q_R-09** | 5'-CAGTGGTGTCAACTCAATTG-3' |
| **C_F-07_RC** | 5'-GTCTTTTACCTCTATCTTTTG-3' | **R** | 5'-TGCACAGTTTTAATTGTG-3' |
| **D** | 5'-AGTGACATAGCAGGAACTACTAG-3' | **S** | 5'-CAATGTATGCCCCTCCCATC-3' |
| **E** | 5'-GAAAAAGGGCTGTTGGAAATG-3' | **S_R-09** | 5'-CAATATATGCCCCTCCTATC-3' |
| **F** | 5'-GGAATTGGAGGTTTTATCAAAG-3' | **T** | 5'-GCAGGAAGCACTATGGGC-3' |
| **G** | 5'-GTAACAGTACTAGATGTGGGTG-3' | **U** | 5'-ATAGAGTTAGGCAGGGATACTC-3' |
| **H** | 5'-GCTGGACTGTCAATGACATAC-3' | **V** | 5'-GAGCCTGTGCCTCTTCAGC-3' |
| **I** | 5'-CCACAGAAAGCATAGTAATAT-3' | **W** | 5'-GTGGCAAGTGGTCAAAAAGTAG-3' |
| **J** | 5'-CACACAAAGGGATTGGAGGAAATGA-3' | **W_R-09** | 5'-GTGGCAAGTGGTCAAAACCCAG-3' |
| **K** | 5'-AATTAGCAGGAAGATGGC-3' | **X** | 5'-GGGACTGGAAGGGCTAATTTAC-3' |
| **K_RC** | 5'-GCCATCTTCCTGCTAATT-3' | **Y_R-09** | 5'-GCTGCTCTTTGCTTGTACTGGG-3' |
| **L** | 5'-CAGCAGTACAAATGGCAGTATTC-3' | | |
